# Neuroendocrine control of synaptic transmission by PHAC-1 in *C. elegans*

**DOI:** 10.1101/2023.08.19.553960

**Authors:** Aikaterini Stratigi, Miguel Soler-García, Mia Krout, Shikha Shukla, Mario De Bono, Janet E. Richmond, Patrick Laurent

## Abstract

A dynamic interplay between synaptic and neuromodulatory signalling guarantees flexible but robust neuronal circuits. Presynaptic modulation plays a crucial role in controlling the excitatory-inhibitory balance within networks. Here, we designed a genetic screen to identify genes involved in the neuromodulation of the *C. elegans* neuromuscular junctions (NMJ) and identified the orthologs of the Protein Phosphatase 1 regulatory subunit PHACTR1 (*phac-1)* and the presynaptic phosphoproteins Synapsin (*snn-1*). Five *de novo* variants of human PHACTR1 are associated with severe early-onset epilepsies (DEE70). To understand the impact of these variants, we introduced the DEE70 mutations into *phac-1*. These mutations resulted in the formation of a constitutively active PP1-PHAC-1 holoenzyme that disrupts cholinergic signalling at the NMJ. By using quantitative fluorescence imaging, electron microscopy and electrophysiology, we found that the constitutive holoenzyme alters the synaptic vesicle cycle, reduces the synaptic vesicle reserve pool, and increases neuropeptide release by dense-core vesicles. Notably, while SNN-1 phosphoregulation contributes to NMJ signalling, genetic interactions suggest that SNN-1 is not the main effector of PP1-PHAC-1 holoenzyme signalling. Collectively, our results confirm the pathogenicity of DEE70 variants, clarify their dominant-positive effects, and provide evidence of a presynaptic mode of action for DEE70.

## Introduction

Neuromodulation by neuropeptides alters circuit dynamics by changing cellular and synaptic properties of specific neurons. The interplay of fast synaptic and slower neuromodulatory communication between neurons ensure circuit adaptability, stability, and the integration of contextual information [1, 2]. Neuropeptides also play a significant role in preserving the balance between excitatory and inhibitory signals within circuits (E/I balance). This homeostatic function of neuropeptides contributes to both fear learning and prevention of pathological disorders such as epilepsy and autism [3–6]

The sinusoidal locomotion of *C. elegans* results from coordinated muscle contraction-relaxation mediated by a set of interconnected cholinergic and GABAergic motor neurons [7, 8]. Mutations producing abnormal activities in the motor circuit can cause aberrant muscle contractions and uncoordinated locomotion [9]. Decades of studies revealed multi-level regulation of the E/I balance in the motor circuit of *C. elegans*. For example, animals carrying a gain of function mutation in the ionotropic acetylcholine (ACh) receptor ACR-2 display spontaneous epileptic-like convulsions. *acr-2(gf)* mutations result in increased endogenous cholinergic excitation and reduced GABAergic inhibition in the locomotor circuit [10, 11]. The activity-regulated production of neuropeptides by the cholinergic motorneurons maintains the E/I balance in the locomotor circuit; e.g. increased production and secretion of the FMRF-like Peptide FLP-18 inhibits convulsions caused by *acr-2(gf)* [12, 13]. The complex homeostatic control of the motor circuit by neuropeptides remains incompletely understood.

Previous genetic screens have not only identified multiple neuropeptides and G-protein coupled receptors (GPCRs) as key modulators of acetylcholine (ACh) and GABA release at the *C. elegans* NMJ [14–16]. Downstream of GPCRs, genetics has highlighted signalling by rho GTPases, protein kinases A (PKA) and C (PKC), and phospholipid pathways to regulate SV dynamics. Among presynaptic effectors of these pathways are phosphoproteins such as UNC-18/STXBP1, UNC-13/Munc-13 or SNN-1/Synapsin [17–24]. The mechanisms identified are conserved, and also modulate presynaptic plasticity in mammalian neurons [25, 26]. Underscoring the importance of presynaptic modulation for circuit homeostasis, pathogenic variants of STXBP1, Unc13A and Synapsin I are associated with human encephalopathies related to E/I imbalance disorders, including epilepsy and autism [27–30].

Here, we designed a genetic screen aimed at understanding the role of FLP-18 signalling in regulating the motor circuit balance. Among its effectors, we isolated the single *C. elegans* orthologs of PHACTR1 (*phac-1*) and Synapsin (*snn-1*). Synapsins are presynaptic phosphoproteins controlling synaptic vesicle (SV) dynamics and the formation of SV clusters [31]. In humans, 4 proteins called the PHosphatase 1 and ACTin Regulators (PHACTR1-4) are Protein Phosphatase 1 family (PP1s) regulatory subunits [32]. The substrates, localisation and functions of PP1s are largely defined by their association with hundreds of regulatory subunits, forming as many specific PP1/protein subunit complexes known as holophosphatases [33]. The binding of PP1 to PHACTR1 profoundly transforms PP1’s surface adjacent to its catalytic site, resulting in improved substrate specificity of the PP1-PHACTR1 holophosphatase [34]. The substrates of the human PP1-PHACTR1 holophosphatase include structural and regulatory cytoskeletal proteins such as Spectrin-a, IRSp53, Ankyrin and Afadin [34]. Several *de novo* variants in PHACTR1 are associated with Developmental Encephalopathy and Epilepsy 70 (DEE70), a syndrome that includes infantile epileptic seizures [35, 36]. DEE are severe forms of early-onset epilepsy caused by rare -often *de novo-* mutations in genes involved in cell migration, proliferation, and organization, neuronal excitability, synaptic transmission, and plasticity [37, 38]. Knockdown in mice suggests that PHACTR1 controls neuronal migration and dendritic tree complexity [36].

Our data demonstrate a role for PHAC-1 and SNN-1 in maintaining the E/I balance of the *C. elegans* motor circuit. We observe that genetic interactions between PHAC-1 and SNN-1 control presynaptic properties. Introducing 3 of the human variants causing DEE70 into the *C. elegans phac-1* gene generates a constitutively active PP1-PHAC-1 holophosphatase. PHAC-1 signalling reduces the SV pool, causes SV recycling defects and increases Dense Core Vesicle (DCV) secretion of neuropeptides. Altogether, our work provides insights into presynaptic modulation mechanisms preventing E/I imbalance disorders.

## Results

### A suppressor screen highlights *snn-1* and *phac-1* involvement in neuromodulation of the neuromuscular junctions

Bioamines and neuropeptides modulate neurotransmitter release at NMJs through synaptic or extra-synaptic GPCR signalling [19, 39–41]. We and others observed that overexpression of the FLP-18 neuropeptide, under its own promoter, alters the locomotion of *C. elegans*: animals reverse more frequently, body bends are deeper during forward locomotion and extremely deep during reversals, leading to slower and uncoordinated locomotion on agarose plates and reduced thrashing frequency in liquid [13, 42, 43] (Supplementary Figure 1). FLP-18 receptors are expressed on sensory neurons, motoneurons and muscles [43, 44]. We took advantage of this phenotype to design a suppressor screen aiming at identifying genes involved in neuropeptide signalling at the worm NMJ (Figure 1a). We first validated this model by crossing the FLP-18 overexpressing strain (*flp-18(XS)*) with 27 mutants defective in genes controlling neuropeptide expression, DCV biogenesis, peptide release, or GPCR signalling. As expected, these mutants modulated the FLP-18-induced uncoordinated locomotion on plates (Figure 1b). Mutants with the most prominent effects also impacted thrashing frequency in liquid (Figure 1d). The strongest suppression was observed for NPR-5, a FLP-18 receptor expressed in sensory neurons and body wall muscles [13, 42, 43]. We next mutagenized the *flp-18* overexpressing strain and isolated 25 strains that were wild-type for their locomotion, despite the FLP-18 overexpression. We identified some of the causal variants by whole genome sequencing and confirmed by other alleles included *npr-5(ulb08), crh-1(ulb06), cmk-1(ulb07), ric-19(ulb12), snn-1(ulb14) and F26H9.2(ulb03),* which we renamed *phac-1* (Figure 1b in blue). The mutations in the CREB transcription factor *crh-1(ulb06)* and the CAMKI *cmk-1(ulb07)* reduced expression from the *flp-18* promoter: our *flp-18* overexpression strain co-expressed GFP in an operon with *flp-18,* and we observed reduced GFP fluorescence. Transcriptional regulation of *flp-18* by *crh-1* was previously reported [44]. Mutations in *ric-19*, the ortholog of Islet Cell Autoantigen (*ICA69),* were previously observed to reduce DCV biogenesis [45]. These data suggest *crh-1, cmk-1* and *ric-19* act in the FLP-18 expressing neurons to promote neuropeptide signalling (Figure 1c). Among the other suppressors identified and confirmed by the thrashing assay, we identified a deletion in the single *C. elegans* ortholog of Synapsin, *snn-1(ulb14),* (Figure 1e). Synapsins are phosphoproteins that tether synaptic vesicles to each other, forming synaptic vesicle clusters. We also identified a deletion in *F26H9.2(ulb03),* the single *C. elegans* ortholog of the 4 mammalian Phosphatase and ACTin Regulators (PHACTRs). The C-terminal domains of mammalian PHACTRs include 3 RPEL motifs (involved in G-actin binding: Pfam PF02755) and one PP1c Binding Domain. These are well conserved in F26H9.2, whereas the unstructured N-terminal region is poorly conserved and the first C-terminal RPEL is only partially conserved (Supplementary Figure 2a). Based on this homology, we renamed F26H9.2 as PP1 Regulatory Protein: *phac-1*. The *phac-1(ulb03)* allele obtained from the screen corresponded to a C-terminal deletion and frameshift removing RPEL motifs and the PP1 binding domain. We obtained two other likely null alleles from the Japanese *C. elegans* consortium that correspond to large C-terminal deletions, *phac-1(tm2463)* and *phac-1(tm2453),* the latter also possessing an early stop codon (Figure 1f, orange).

**Figure 1.**
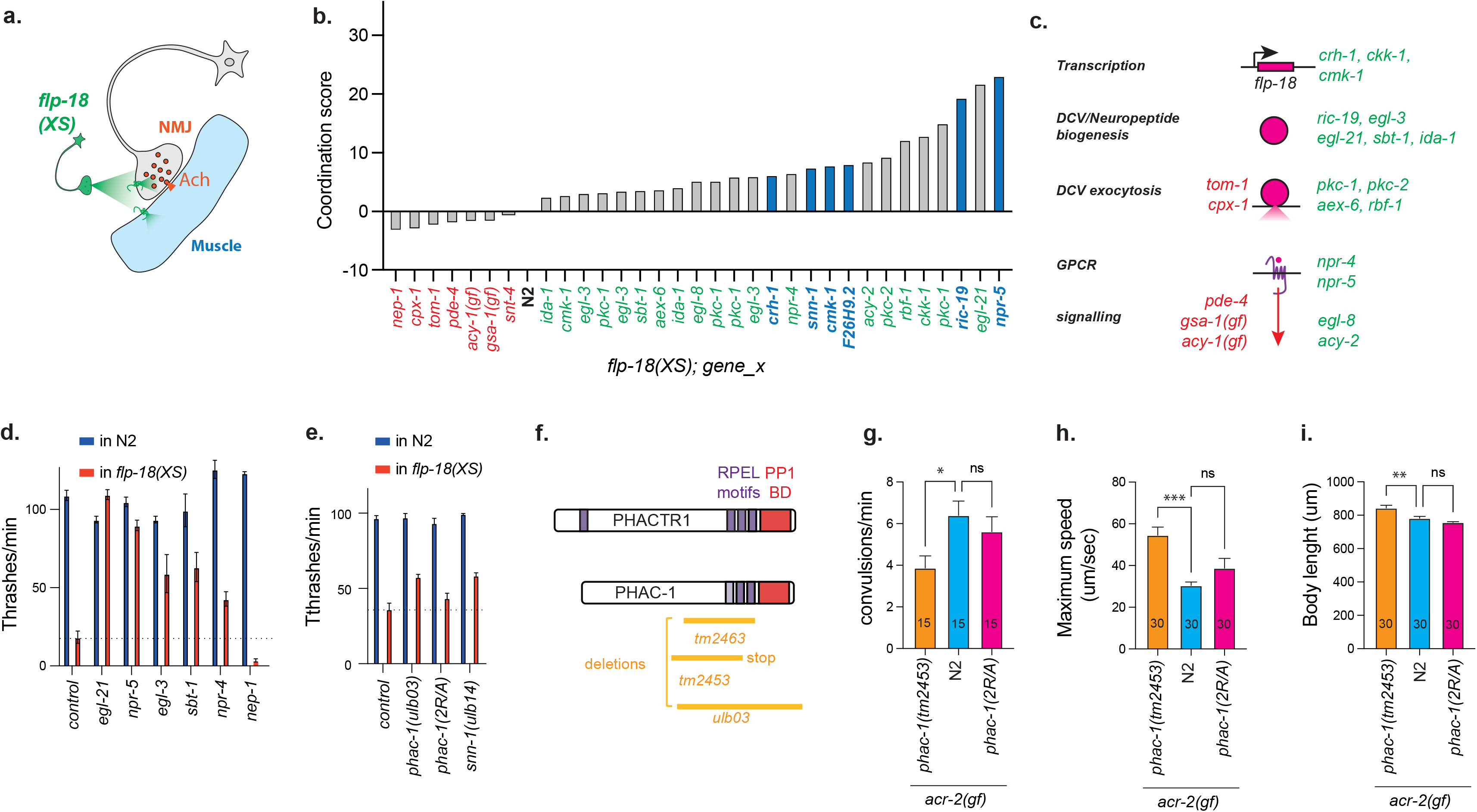
A suppressor screen allows the identification of genes involved in neuromodulation of the NMJ by FLP-18. **a)** Overexpression of the neuropeptide FLP-18 (*flp-18(XS)*) alters the locomotion of *C. elegans* though modulation of the NMJ. This phenotype was used to identify genes reducing neuropeptide signalling at the NMJ. The screen uncovered genes involved in neuropeptide transcription and biogenesis, DCV exocytosis, and neuropeptide signalling at the NMJ. **b)** Locomotion was scored for the FLP-18 overexpression strain crossed with mutants expected to either enhance (in red) or reduce (in green) FLP-18 function, relative to the N2 wildtype background (black). Strains isolated from the FLP-18 suppressor screen are highlighted in blue; the causal genes identified include *npr-5, crh-1, cmk-1, ric-19, snn-1* and F16H9.2/*phac-1*. **c)** The control genes were selected across genes involved in neuropeptide and DCV biogenesis, secretion and signalling. **d)** The frequency of thrashes in liquid was quantified for each mutant in the N2 genetic background (in blue) and in the *flp-18(XS)* background (in red). The frequency of thrashes is strongly reduced by FLP-18 overexpression. The strongest suppressors of the *flp-18(XS*) thrashing phenotype are *egl-21* and *npr-5*. **e)** The frequency of thrashes in liquid for a subset of suppressor mutants identified in the forward genetic screen. *phac-1(ulb03)* and *snn-1(ulb14)* partially suppress the *flp-18(XS)* thrashing phenotype. **f)** Schematic drawing of the structure of PHACTR1 and PHAC-1. Green: RPEL motifs. Orange: PP1-binding domain. The deletions corresponding to the null alleles assayed are represented below in orange. **g)** Quantification of convulsion frequency in genotypes as indicated. The convulsion rate is reduced in *acr-2(gf);phac-1(tm2453)* compared to *acr-2(gf).* **h)** The maximum forward speed of locomotion is increased in *acr-2(gf);phac-1(tm2453)* compared to *acr-2(gf).* **i)** The body length is increased in *acr-2(gf);phac-1(tm2453)* compared to *acr-2(gf).* Data on 1d, 1e,1g, 1h, 1i are plotted as the means ± SEM. Statistics use One-way ANOVA and *Tukey’s* multiple comparison test. *:P≤0.05, **:P≤0.01, ***:P≤0.001.

Under normal cultivation conditions, the *phac-1* deletion mutants grew and developed as well as wild-type controls (N2). However, their speed of locomotion was reduced (Supplementary Figure 2b). Neuroendocrine regulation of the motor circuit by *flp-18* was previously shown to reduce convulsions of an *acr-2(gf)* strain [13]. We crossed *phac-1* mutants into the *acr-2(gf)* strain to test the potential role of *phac-1(tm2453)* in the control of the E/I balance of the motor circuit. We observed that *phac-1(tm2453)* reduced the convulsions induced by the *acr-2(gf)* allele (Figure 1g). *phac-1(tm2453)* also improved the slow locomotion and the hypercontracted phenotype associated with the *acr-2(gf)* mutation (Figure 1h, 1i). As we suspected alterations in NMJ neurotransmission in the *phac-1* deletion mutants, we used the Aldicarb-induced paralysis assay to determine whether NMJ signalling was altered in freely moving worms. ACh and GABA promote contraction and relaxation of body-wall muscle, respectively. Application of the acetylcholinesterase inhibitor Aldicarb prevents ACh hydrolysis in the synaptic cleft. The resulting accumulation of ACh causes hypercontracted paralysis within hours. Mutants that reduce ACh release, increase GABA release, or alter ACh / GABA signalling in the body wall muscle, confer resistance to Aldicarb paralysis. Conversely, Aldicarb-hypersensitivity occurs in mutants that increase ACh signalling or decrease GABA signalling at the NMJ [46] (Figure 2a). For all Aldicarb assays, we measured the full-time course for paralysis in the presence of Aldicarb and determined the time point at which 50% of animals are paralysed for 3 to 7 independent assays, corresponding altogether to 60 to 140 animals. All *phac-1(tm2453), phac-1(ulb03)* and *phac-1(tm2463)* deletion mutants caused similar Aldicarb-hypersensitive phenotypes, suggesting a higher rate of ACh release or altered E/I balance (Figure 2b).

**Figure 2.**
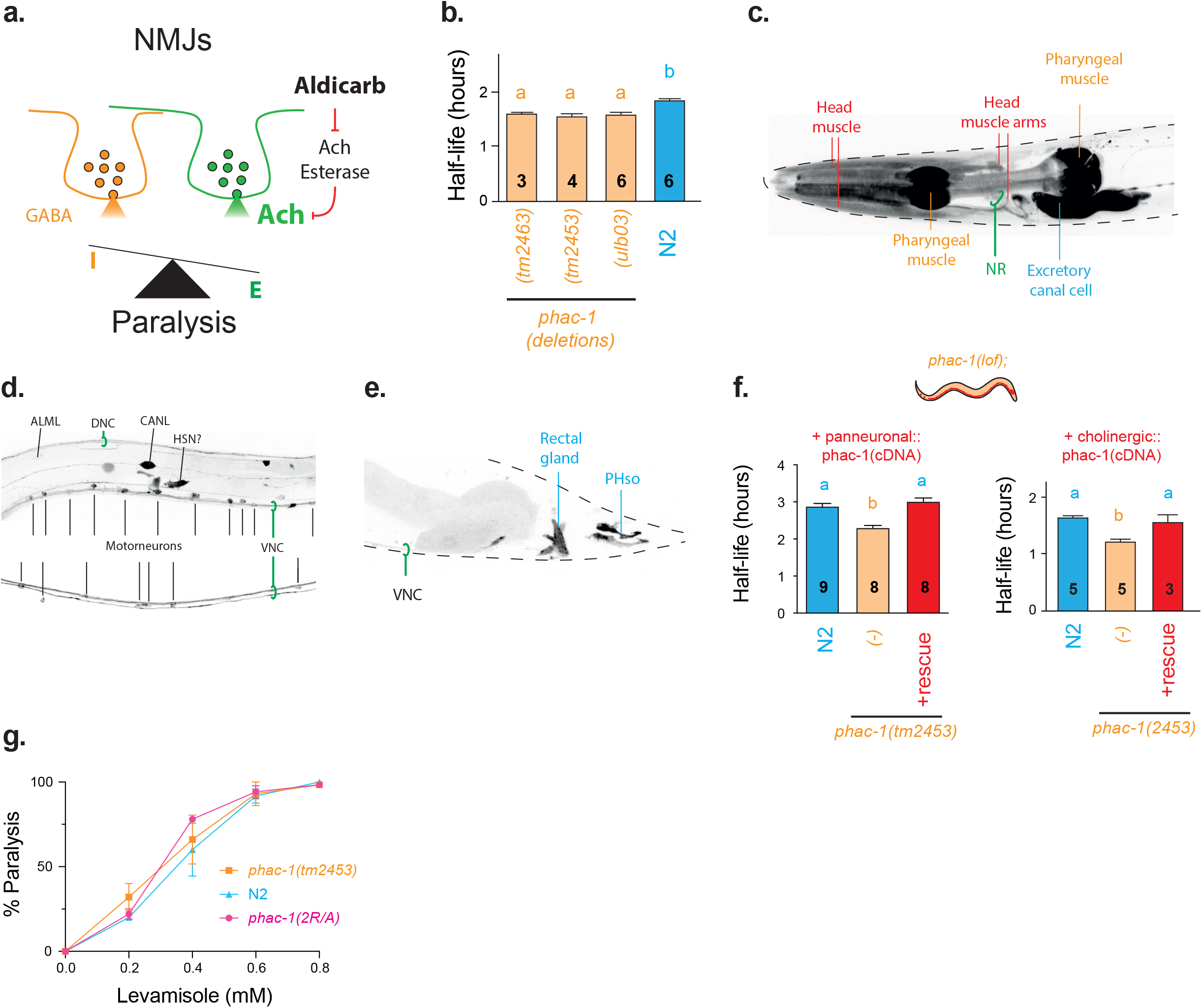
The *C. elegans* ortholog of PHACTRs regulates neurotransmission at cholinergic NMJ. **a)** The Aldicarb assay was used to detect modulation of neurotransmission at the NMJ. Aldicarb inhibits acetylcholinesterase, leading to accumulation of Ach, which produces paralysis. The rate of paralysis reflects the strength of ACh and GABA signalling at the NMJ. We determined the time point at which 50% of animals are paralysed by plotting data obtained from 20-30 animals in several independent assays (Ns are indicated in the histograms). **b)** All *phac-1* deletion mutants are Aldicarb-hypersensitive, suggesting higher Ach or lower GABA signal than controls. The number of Aldicarb assays is indicated for each condition. c-e) The adult expression pattern of TagRFP was determined in a reporter strain carrying *Ex[phac-1-SL2TagRFP]* (see Supplementary Figure 2). **c)** In the head, expression is observed in the pharynx, muscle, excretory cells and nerve ring (NR). **d)** In the midbody, expression is observed in motor neurons of the ventral nerve cord (VNC), and other neurons as labelled. Images from two worms fed with a *phac-1* RNAi 24hr prior to imaging are shown. **e)** In the tail, expression is observed in neurons, phasmid socket cells (PHso) and in a cell identified as the rectal gland. **f)** The Aldicarb-hypersensitivity of *phac-1* null mutants is rescued by the expression of *phac-1* cDNA in all neurons using the *rab-3* promoter (left) or only in cholinergic neurons using the *unc-17* promoter (right). **g)** Sensitivity to the ionotropic cholinergic agonist levamisole is not altered in *phac-1* mutants. The *phac-1(2R/A)* is described later in text. Data on 2b, 2f and 2g are plotted as the means ± SEM. Significant differences in pairwise comparisons are indicated by compact letters display. In short, genotypes that share the same letter are statistically indistinguishable to each other, genotypes that have a different letter statistically differ (P<0.05) using One-way ANOVA and *Tukey’s* multiple comparison test.

### *phac-1* is expressed broadly and regulates neurotransmission at the cholinergic NMJ

To determine the expression pattern of *phac-1,* we generated a knock-in strain in which wrmScarlet was fused to the C-terminal of PHAC-1. In this reporter strain, wrmScarlet was weakly but broadly expressed. We could identify expression in pharyngeal muscles, head muscles, excretory cells, and unidentified neurons that project to the nerve ring (Supplementary Figure 2c). To better observe neuronal expression patterns, we generated an overexpression reporter strain in which a 6kb endogenous locus, including 3kb of promoter sequence, drives the expression of TagRFP. TagRFP is inserted downstream of the stop codon of *phac-1*, forming an operon. The expression pattern appeared similar, including strong expression in pharyngeal muscle, head muscle cells, excretory gland and canal cells, phasmid and amphid socket glia and neurons projecting to the nerve ring (Figure 2c, 2d, 2e). TagRFP was expressed from the early embryo to the adult stage (Supplementary Figure 2d). To better define the neuronal pattern of expression, this strain was fed with an RNAi clone to knock-down *phac-1* and TagRFP [47]. As neurons are relatively resistant to RNAi knockdown, we could observe expression of TagRFP in cholinergic motoneurons of the ventral nerve cord as well as in AVL, AVM, CAN, HSN among other neurons (Figure 2d). N and C-terminal GFP fusion constructs expressed in neurons localised to the cytoplasm and neuritis, but not the nucleus (Supplementary Figure 2e). As *phac-1* was broadly expressed, we tested whether neuronal expression could rescue the Aldicarb-hypersensitive phenotype. The Aldicarb response was fully restored to *phac-1(tm2453)* mutants that express *phac-1* cDNA panneuronally or selectively in cholinergic neurons (Figure 2f). The muscle sensitivity to the ionotropic cholinergic agonist levamisole was not significantly altered in *phac-1* deletion mutants (Figure 2g). These results confirm that Aldicarb-hypersensitivity in *phac-1(tm2453)* mutants correspond to abnormal presynaptic signalling rather than a defect in muscle response to ACh neurosecretion.

### Mutations in *phac-1* that mimic the human DEE70 variants generate a constitutively active holophosphatase

To evaluate the conservation of PHACTRs function, we attempted to rescue *phac-1(tm2453)* with the human gene. Panneuronal expression of human PHACTR1 cDNA rescued the Aldicarb-hypersensitivity of *phac-1(tm2453),* suggesting that at least some PHACTR functions are conserved in neurons (Figure 3a). Human genetic studies previously revealed 4 heterozygous dominant *de novo* mutations in PHACTR1 associated with DEE70. These 4 mutations correspond to substitutions of highly conserved amino acids located within the last 2 RPEL motifs [35, 36]. These RPEL motifs are adjacent to the PP1-binding site, resulting in steric hindrance between G-actin and PP1 for binding to PHACTRs. As a result, it is believed that PHACTRs cycle between an active state as PP1-PHACTRs holophosphatase and an inactive state as PHACTR-G-actin, depending on the local concentration of G-actin [48–50]. We introduced 3 of these DEE70 variants, *L500P*, *L519R*, and *R521C,* in the *C. elegans phac-1* gene using CRISPR-Cas9, substituting the corresponding amino acids in PHAC-1 (Figure 3b). In contrast to *phac-1* deletion mutants, which caused Aldicarb-hypersensitivity, the 3 alleles mimicking human *DEE70* alleles caused Aldicarb-resistance (Figure 3c, 3d). Other mutations in highly conserved Arginines (R) are also known to disrupt PHACTR1/4 RPEL motifs, which prevent G-actin binding and induce the constitutive binding of PP1 [48, 49]. We generated a *phac-1*(2R/A) allele where the R of the last 2 **R**PEL motifs of *phac-1* were replaced by Alanines (A) as done in [48, 50]. We compared the effects of 2R/A mutations to the DEE70 mutations. Similar to the 3 *phac-1* alleles mimicking human *DEE70*, the *phac-1*(2R/A) allele caused Aldicarb-resistance (Figure 3c, 3d). Conversely, an *R536P* mutation replicating an hypomorphic allele of PHACTR4 [51], caused a weak but significant Aldicarb-hypersensitivity (figure 3d). Therefore, *DEE70* mutations and 2R/A mutations caused a *gain-of-function* (*gf*) phenotype, likely by inducing constitutively active PP1-PHAC-1 holophosphatase. To confirm this, we expressed cDNA corresponding to the *phac-1*(2R/A) allele in the N2 background. The expression of *phac-1*(2R/A) *cDNA* in all neurons, or in cholinergic motoneurons, caused Aldicarb-resistance, suggesting a dominant-positive mode of action in cholinergic motoneurons (Figure 3e). Animals heterozygous for *phac-1(2R/A)* and *phac-1(L500P)* were also Aldicarb-resistant compared to N2 controls (Figure 3f). Hence, *DEE70* mutations act as dominant positive alleles at adult cholinergic NMJ to produce Aldicarb-resistance. For simplicity, in the following sections, we refer to *phac-1(tm245*3) as *phac-1(lf)* and *phac-1(2R/A)* as *phac-1(gf).* To gain support for the interpretation that *DEE70* and 2R/A mutations cause Aldicarb-resistance by promoting the constitutive binding of PP1 to PHAC-1, we generated a protein fusion in which PHAC-1 is constitutively fused to GSP-1, one of the 4 Protein Phosphatase 1 catalytic subunits of *C. elegans*. Panneuronal expression of this fusion protein in *phac-1(lf)* induced Aldicarb-resistance, as observed in *phac-1(DEE70)* alleles and in *phac-1(gf)* (Figure 3g). A D64A mutation within the PP1 catalytic site reduces its enzymatic activity [52]. The panneuronal expression of a mutated PHAC-1-GSP-1(D64A) construct in *phac-1(lf)* had no effect, suggesting PP1 enzymatic activity is required for the fusion protein to confer Aldicarb-resistance. As the *DEE70* and 2R/A alleles mimic the Aldicarb-resistance observed by PP1-PHAC-1 fusion, we conclude the *DEE70* and 2R/A alleles generate PHAC-1 variants that bind PP1 constitutively.

**Figure 3.**
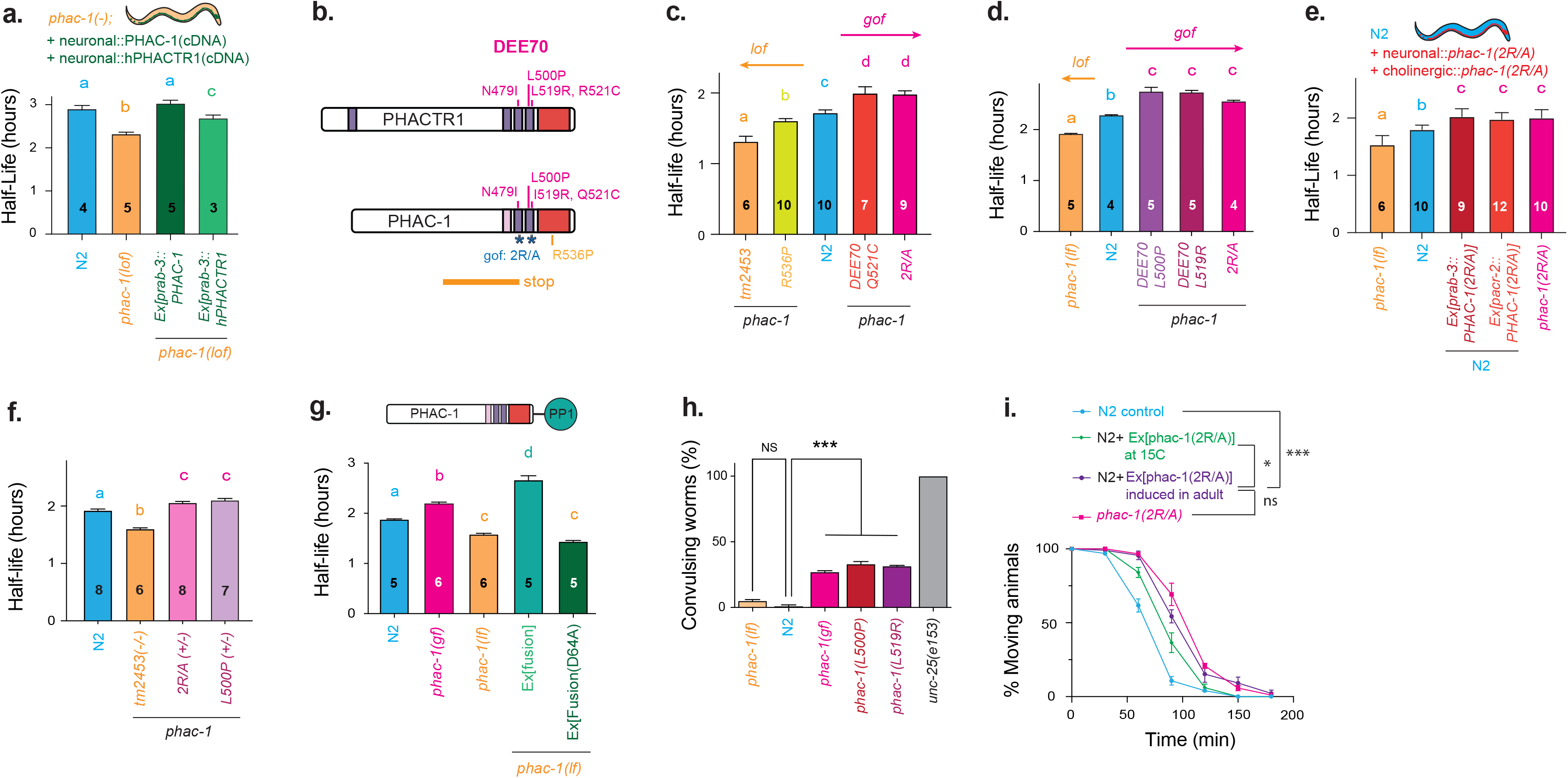
Mutations in *phac-1* that mimic the human DEE70 variants generate a constitutively active holophosphatase. **a)** Pan-neuronal expression of human PHACTR1 cDNA rescues the Aldicarb-hypersensitivity of the *phac-1(lf)* mutant, suggesting conservation of some of the neuronal functions of the protein. **b)** The locations of DEE70 variants of PHACTR1 (top) and the corresponding mutations generated by CRISPR in PHAC-1 are indicated in magenta (bottom). The *2R/A* mutations correspond to *R469A* combined with *R507A*. All labels for point mutations made in *C. elegans* corresponds to the amino-acid numbering of the human protein. The *R536P* mutation corresponds to the *humdy* hypomorph mutant in the mouse PHACTR4 [51]. **c, d)** In contrast to the *phac-1(tm2453)* deletion mutant, the G-actin-binding deficient mutant *phac-1(2R/A)* causes Aldicarb-resistance. **c)** The *Q521C* mutation mimicking the R521C DEE70 also causes Aldicarb-resistance, while the humdy/*R536P* mutation causes a weak Aldicarb-hypersensitivity. **d)** The *L500P* and *L519R* mutations mimicking DEE70 cause Aldicarb-resistance. **e)** Panneuronal or cholinergic expression of a *phac-1(2R/A)* cDNA in the N2 background cause Aldicarb-resistance, suggesting a dominant-positive mode of action for the mutations. **f)** Heterozygous animals carrying *2R/A* or *L500P* mutations are resistant to Aldicarb compared to N2, suggesting a dominant mode of action. **g)** Top, the PHAC-1-PP1 fusion protein in which PP1 corresponds to the GSP-1 subunit of PP1. Panneuronal expression of a PHAC-1-GSP-1 fusion protein in *phac-1(lf)* phenocopies *phac-1(gf)*. Panneuronal expression of the same fusion protein construct carrying a D64A mutation that reduces GSP-1 enzymatic activity has no effect. **h)** In the presence of PTZ, a GABA receptor antagonist, E/I imbalance in the motor network produce neck convulsions. We determined the percentage of worms convulsing for ∼120 animals in 2 independent assays. *phac-1(tm2453)* mutants did no display convulsions while phac-1(gf), *phac-1(L500P)* and *phac-1(L519R)* are PTZ-hypersensitive, as the *unc-25(e156)* controls. **i)** Expression of *phac-1(2R/A)* cDNA 16 hours before adulthood is sufficient to induce Aldicarb-resistance, suggesting *phac-1(2R/A)* impacts neurotransmission in the adult NMJ, post-developmentally. One-way ANOVA, *Tukey’s* multiple comparison test. *:P≤0.05, **:P≤0.01, ***:P≤0.001. All data are plotted as the mean ± SEM. For a, c, d, e, f, g, significant differences in pairwise comparisons are indicated by compact letters display. In short, genotypes that share the same letter are statistically indistinguishable to each other, genotypes that have a different letter statistically differ (P<0.05) using One-way ANOVA and *Tukey’s* multiple comparison test.

DEE70 disorder is characterized by severe early-onset seizures. Pentylenetetrazole (PTZ) is a GABAR antagonist that induces seizures in mammals, and convulsions in *C. elegans* mutants of GABAergic signalling [53]. To test whether GABAergic signalling is reduced in *phac-1* mutants, worms were exposed to PTZ treated plates. We noted a higher occurrence of convulsions in *phac-1(gf), phac-1(L500P),* and *phac-1(L519R)* variants compared to the control N2, while *phac-1(lf)* mutants did not show a similar increase (Figure 3h). PHAC-1 regulation of neurosecretion could reflect changes in neurodevelopment, neuronal or synaptic function. To determine the temporal requirements of *phac-1,* we induced *phac-1(2R/A)* expression at L1 larval stage or adulthood. The induction of *phac-1(2R/A)* in N2 adults induced Aldicarb-resistance, while expression in early larval stage did not (Figure 3i). These results suggest *phac-1(2R/A)* acts post-developmentally in the adult to regulate neurosecretion.

### *phac-1* signalling alters presynaptic pools of synaptic vesicles (SVs) and dense core vesicles (DCVs)

To determine the effects of *phac-1* on the cholinergic motoneurons, we used a combination of quantitative fluorescence imaging, electron microscopy and electrophysiology. We first determined the effects of *phac-1* on the cell biology of the DA cholinergic motoneurons using a panel of strains expressing integrated presynaptic markers [54, 55]. The dorsal axons of the DA motoneurons form stereotypical *en-passant* NMJs. The fluorescence intensity and distribution along these dorsal axons were analysed for each fluorescently tagged protein (Figure 4a, 4b). We did not observe significant differences for the active zone markers UNC-10-GFP suggesting that neither the density of active zones nor their organisation are modified in *phac-1* mutants (Supplementary Figure 3a, 3b, 3c). To observe presynaptic SV pools we used the SV-associated RAB-3-GFP marker. The fluorescence intensity of the RAB-3-GFP peak was reduced in *phac-1(gf)* but not affected in *phac-1(lf)* (Figure 4c). To observe the DCV pool, we used the NLP-21-GFP neuropeptide marker. We observed reduced NLP-21-GFP fluorescence intensity in the axons of *phac-1(gf)* mutants, but no change in soma fluorescence (Figure 4d, Supplementary Figure 3d, 3e). As a proxy to quantify NLP-21-GFP neuropeptide release, we measured the GFP fluorescence intensity in coelomocytes that are known to scavenge NLP-21-GFP released by DA neurons into the pseudocoelomic cavity (Figure 4e) [55]. We observed an increased in coelomocyte NLP-21-GFP fluorescence intensity in *phac-1(gf)* (Figure 4f). These data suggest that increased PHAC-1 signalling reduces the SV pool and increases the release of neuropeptides from DCVs located in the axon of DA cholinergic motoneurons.

**Figure 4.**
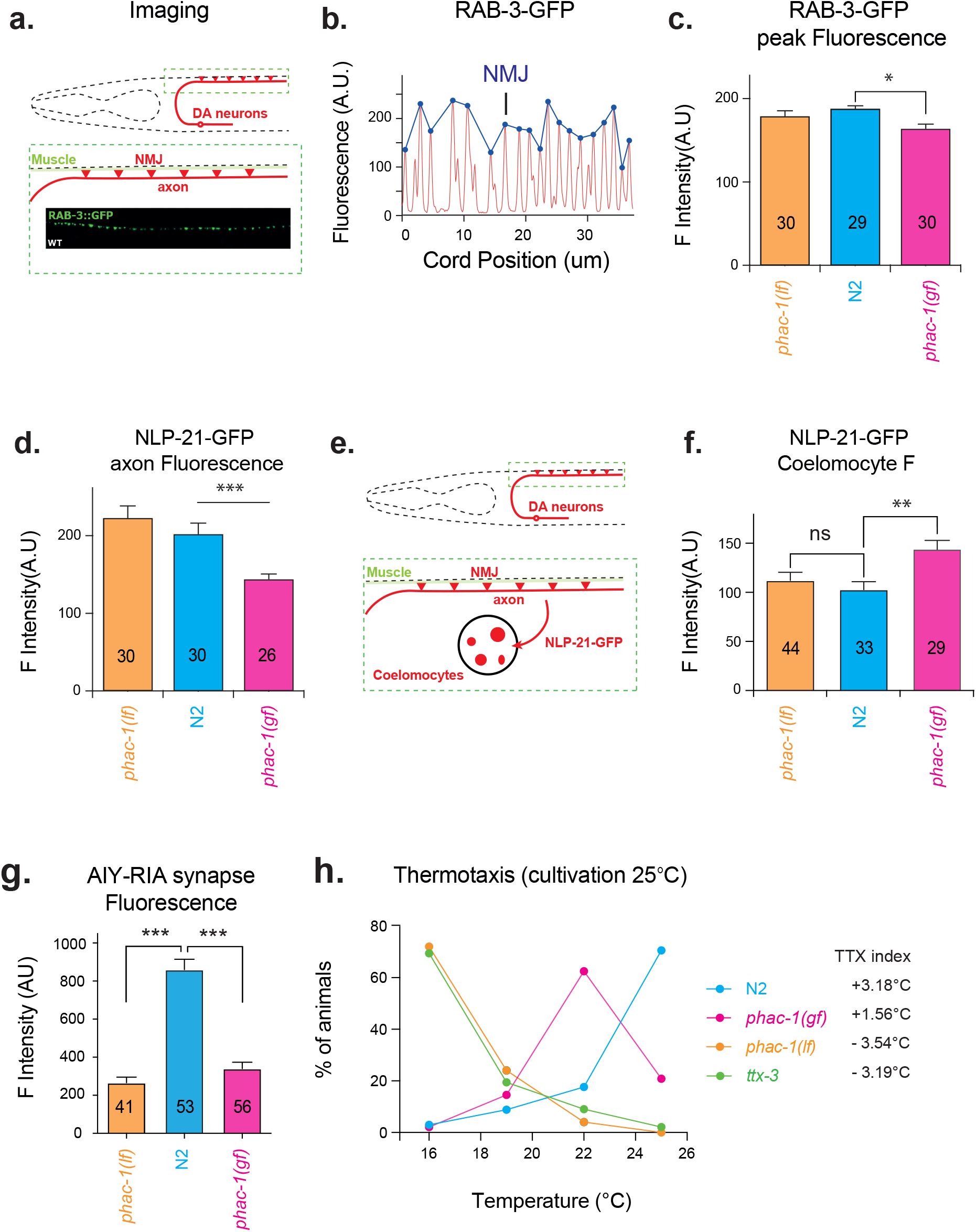
*phac-1* signalling alters the presynaptic pools of synaptic vesicles and dense core vesicles. **a)** Schematic representation of the NMJs formed by the DA cholinergic motoneurons along the dorsal cord, where imaging of synaptic markers was performed. **b)** Representative graph showing fluorescence of the synaptic vesicle (SV) pool marker, RAB-3 along the dorsal nerve cord. Fluorescence peaks correspond to the enrichment of RAB-3 at presynaptic puncta. **c)** The fluorescence intensity at RAB-3-GFP peaks (see method for peak detection) is significantly reduced in *phac-1(gf)* compared to N2, indicating a reduced SV pool. The z-test was used to determine significance values relative to N2 controls. **d)** The fluorescence intensity of the neuropeptide/dense core vesicle marker NLP-21 along the DA axon is significantly reduced in *phac-1(gf)* compared to N2, indicating reduced neuropeptide/DCV biogenesis and/or increased neuropeptide secretion. **e)** Coelomocytes cells scavenge NLP-21-GFP released in the pseudocoelomic cavity; NLP-21-GFP fluorescence in coelomocytes is a proxy for NLP-21-GFP release. **f)** The fluorescence intensity of NLP-21-GFP in coelomocytes is increased in *phac-1(gf)* compared to N2, suggesting increased neuropeptide release. **g)** The fluorescence intensity of the SV marker RAB-3-GFP is reduced in the AIY-RIA synapse of *phac-1(lf)* and *(gf)* mutants compared to N2, suggesting a reduced number of SV at this synapse. **h)** Thermotaxis behaviour of N2, *ttx-3* and *phac-1 (lf)* and *(gf)* mutants habituated to 25°C on food for 24 hrs prior to the assay. *phac-1(lf)* animals show cryophilic behaviour whereas *phac-1(gf)* animals show an intermediate behaviour. Thermotaxis indexes are indicated. Data on 4c, 4d, 4f, 4g are plotted as the mean ± SEM. Statistics use One-way ANOVA and *Tukey’s* multiple comparison test. *:P≤0.05, **:P≤0.01, ***:P≤0.001.

To determine whether the synaptic defects observed at NMJs in *phac-1* mutants were also observed at other synapses, we examined the organisation and function of the synapse formed between AIY and RIA interneurons, two key components of the thermotaxis circuit [56, 57]. Interestingly, the synapse between AIY and RIA displayed reduced RAB-3-GFP fluorescence in both *phac-1(gf)* and *phac-1(lf)* (Figure 4g). *C. elegans* raised on food at 16°C or at 24°C display thermotaxis toward these food-associated temperatures [58]. When cultivated at 16°C in the presence of food, *phac-1(lf)* and *phac-1(gf)* displayed normal cryophilic behaviour (Supplementary Figure 3f). Interestingly, when cultivated at 24°C in the presence of food, we observed defective thermotaxis toward high temperature in *phac-1* mutants (Figure 4h). Such cryophilic behaviour was previously observed in *ttx-3*, which affects the development of the AIY interneurons [56]. However, apart from the RAB-3-GFP fluorescence intensity in presynapses, we did not observe morphological differences for AIY in *phac-1* mutants (Supplementary Figure 3g), suggesting the function rather than the development of the thermotaxis circuit is affected in *phac-1* mutants.

### The synaptic vesicle cycle is altered in *phac-1* mutants

*C. elegans* NMJs exhibit graded synaptic ACh and GABA transmission, and endogenous neural activity continuously drives neurotransmitter release [59]. To record stimulus-evoked ACh release, we applied a 20V electric stimulus to the ventral nerve cord evoking Ach release at the NMJ [18, 60]. Under these conditions, *phac-1(lf)* and *phac-1(gf)* mutants displayed the same evoked EPSC amplitude, charge transfer, and kinetics as the N2 controls (Figure 5a-d). These results suggest that exocytic properties of cholinergic motorneurons are not modified in *phac-1* mutants in these recording conditions. To better define the synaptic defects that cause the changes in presynaptic pools of SV and neuropeptide/DCVs, we used transmission electron microscopy (TEM) after high-pressure freeze fixation. We confirmed the ultrastructural integrity of the ventral cholinergic NMJs in *phac-1* mutants, but observed smaller dense projections in *phac-1(gf)* compared to N2 and *phac-1(lf)* (Supplementary Figure 4a-c). We observed larger presynaptic terminals in *phac-1(lf),* a phenotype thought to result from defects in SV recycling as terminal size is maintained by a balance in exo- and endocytosis (Figures 5e) [61, 62]. We also observed a marked reduction in the total number of SVs in *phac-1(gf)* (Figure 5f). Given the reduced cholinergic signalling in *phac-1(gf),* this suggests a defect in SV biogenesis and/or SV recycling. We also observed an increase in average SV diameter in *phac-1(lf)* and *(gf)*, a phenotype observed in endocytic mutants and suggestive of alterations in the SV recycling/sorting process (Figure 5g) [62–64]. A slight increase was observed in *phac-1(lf)* for vesicles docked on the plasma membrane adjacent to dense projections (DPs) that likely represent primed, fusion-competent SVs (Figure 5h). Together, these EM observations suggest that the available SV pool in *phac-1(gf)* mutants is reduced, possibly because of impaired SV recycling, while more fusion-competent SVs were observed in *phac-1(lf).* We also observed more DCVs in the synaptic profiles of *phac-1(lf)* mutants when compared to controls and *phac-1(gf)* (Figure 5i). These DCVs were clustered in distal regions of the synaptic terminals relative to the DP, matching the increased density of NLP-21-GFP peaks in the axon of *phac-1(lf)* axons (Figure 5j, 5k, respectively). To further explore a potential imbalance of SV exocytosis and endocytosis in these *phac-1* mutants, we used the pH-sensitive SNB-1-pHluorin marker expressed in cholinergic motoneurons. The membrane protein Synaptobrevin/SNB-1-pHluorin cycles between the SV membrane and the plasma membrane during the SV cycle. As SNB-1-pHluorin fluorescence is quenched in the acidic environment of the recycled SV lumen, the fluorescent signal corresponds to SNB-1-pHluorin present in the plasma membrane following exocytosis, when pHluorin is exposed to the extracellular milieu at neutral pH. Thus, SNB-1-pHluorin fluorescence can reflect the balance between SV exocytosis and endocytic rates (Figure 5l) [65]. We observed increased SNB-1-pHluorin fluorescence in dorsal cholinergic NMJs of both *phac-1(lf)* and *phac-1(gf)* mutants (Figure 5m). In contrast to SNB-1-pHluorin fluorescence, total SNB-1-GFP fluorescence intensity was not significantly different between either *phac-1* mutants and control animals (Supplementary Figure 4d, 4e). Finally, we quantified the recruitment of the endocytic markers ITSN-1-GFP and UNC-57-RFP at the presynaptic terminals of *phac-1* compared to controls. We observed reduced ITSN-1-GFP and UNC-57-RFP in *phac-1(gf),* compared to controls and *phac-1(lf)*, which would be indicative of endocytic defects (Figure 5n, 5o, respectively). The combination of increased SNB-1-pHluorin fluorescence, reduced ITSN-1-GFP and UNC-57-RFP levels, reinforce the EM data suggesting that *phac-1(gf)* mutants have endocytic defects. Thus, we conclude that *phac-1* signalling regulates SV recycling at *C. elegans* cholinergic NMJs.

**Figure 5:**
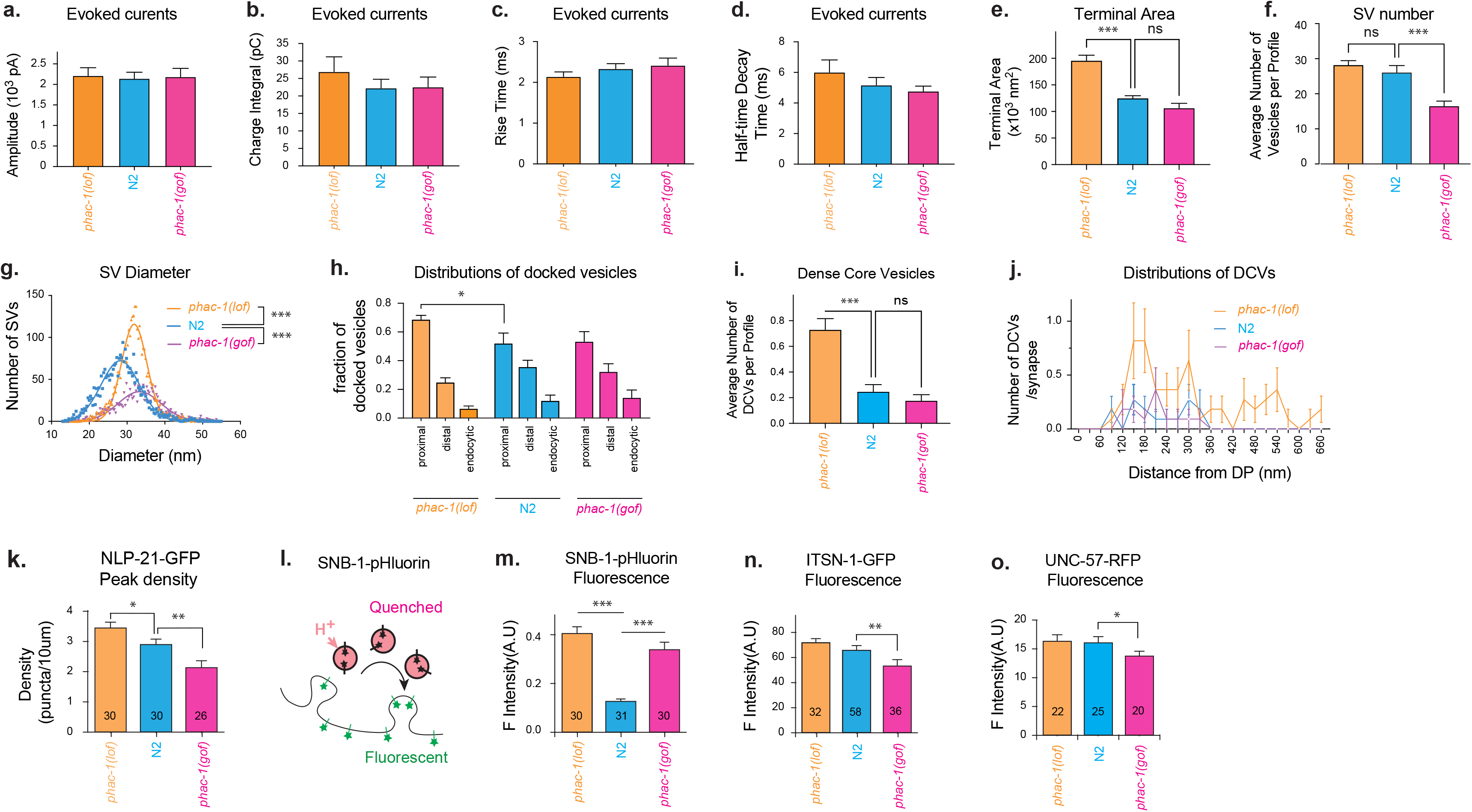
Despite normal excitability and exocytic properties of cholinergic neurons, the synaptic vesicle cycle is altered in *phac-1* mutants. **a-c)** Excitatory postsynaptic currents (EPSC) at the NMJ of N2 and *phac-1(lf) and (gf)* mutants were recorded after a single 20V pulse onto the ventral nerve cord (Ringer + 5mM Ca^2+^). The quantification of the evoked currents does not reveal differences in EPSC amplitude **(a)**, charge transfer **(b)**, rise time **(c)** and half-time decay **(d)** in *phac-1* mutants compared to N2. All data are expressed as mean ± SEM. The Mann Whitney T-test was used to determine significance values relative to N2 controls. **e)** The area of the presynaptic terminals is increased in *phac-1(lf)* compared to N2, suggesting imbalance between exocytosis and endocytosis in *phac-1(lf)* neurons. **f)** The average number of synaptic vesicles (SV) per TEM profile containing a DP shows a strong reduction in *phac-1(gf) mutants.* g) The mean diameter of the SVs was larger in *phac-1(lf)* and *(gf)* mutants compared to N2 (Gaussian non-linear fit: 31.8, 33.3 and 28.2 nm diameter, respectively) suggestive of SV recycling defects. **h)** The fraction of docked SVs in the proximal (0-90nm), distal (91-300nm) and the endocytic zone (301-500nm) relative to the DP indicates a higher proportion of SVs primed next to the DP in *phac-1(lf)*, suggesting higher release probability. **i)** The average number of DCVs per TEM profile is increased in *phac-1(lf)* compared to N2. **j)** The distribution of DCVs along the axon relative to DP suggesting accumulation of DCVs away from the presynaptic terminals in *phac-1(lf)*. **k)** Clusters of DCV carying NLP-21-GFP are observed along the DA axon as fluorescence peaks (see method for peak detection). The peak density is increased in *phac-1(lf)* and decreased in *phac-1(gf)* compared to N2. **l)** The pH-sensitive marker SNB-1-pHluorin is quenched in the acidic environment of the SVs. Its fluorescence reports the fraction of SNB-1 at the plasma membrane and reflects the balance between SV exocytosis and endocytosis. **m)** The fluorescence intensity of SNB-1-pHluorin in the dorsal nerve cord is increased in *phac-1(lf)* and in *phac-1(gf)* compared to N2. **n)** The fluorescence intensity of the endocytic marker ITSN-1-GFP is reduced in the dorsal nerve cord of *phac-1(gf)*, suggesting a defect in endocytosis. **o)** The fluorescence intensity of the endocytic marker UNC-57-RFP is reduced in the dorsal nerve cord of *phac-1(gf)*, suggesting a defect in endocytosis. Data are plotted as the mean ± SEM. Statistics use One-way ANOVA and *Tukey’s* multiple comparison test. *:P≤0.05, **:P≤0.01, ***:P≤0.001.

### *phac-1* and *snn-1* act together to regulate the NMJ

As *snn-1(ulb14)* was identified together with *phac-1(ulb03)* as suppressors of the FLP-18 overexpression phenotype, we further dissected the genetic interaction between *phac-1(lf)*, *phac-1(gf)* alleles, and the *snn-1(tm2557)* null allele previously described in [24]. First, we observed that the Aldicarb-resistance of *phac-1(gf)* mutants was strongly reduced by the *snn-1(tm2557)* mutation, whereas the Aldicarb-hypersensitivity of *phac-1(lf)* was weakly reduced by *snn-1(tm2557)* (Figure 6a). These results fit a model in which Synapsin/*snn-1* might mediate the Aldicarb-phenotypes of both *phac-1(gf)* and *phac-1(lf)*. Interestingly, Serine 9 of Synapsin I appeared amongst the candidate peptide substrates of PP1-PHACTR1 holophosphatase identified by a SILAC approach in mouse primary neurons [34]. To directly analyse the phosphorylation state of Serine 9 SNN-1, we designed antibodies against a 13 amino acid peptide that included phosphorylated serine 9 of SNN-1 [24]. Analysis of SNN-1 by Western blot revealed that phosphorylation of serine 9 was increased in *phac-1(lf)* and decreased in *phac-1(gf)* (Figure 6b, Supplementary Figure 4i). The specificity of the phosphoantibody was confirmed using a null mutant strain for SNN-1 and a strain in which Serine 9 of SNN-1 was replaced by an Alanine. Weak binding to SNN-1(S9A) was observed suggesting residual binding affinity to dephospho-SNN-1. The hydrophobic N-terminal domain of Synapsin binds with high affinity to SV phospholipids and phosphoregulation of serine 9 controls its ability to bind SVs, thereby altering SV dynamics [31, 66, 67]. We tested the Aldicarb response of phosphodeficient *snn-1(S9A),* alleles where the phosphorylated serine is replaced by an alanine [24]. If the dephosphorylation of serine 9 mediates the Aldicarb resistance of *phac-1(gf)*, the phosphodeficient allele of *snn-1(S9A)* should mimic the Aldicarb resistance of *phac-1(gf)* and supress the Aldicarb-hypersensitivity of *phac-1(lf).* Although the *snn-1(S9A)* allele replicated the Aldicarb-resistance of *phac-1(gf)*, it did not supress the Aldicarb-hypersensitivity of *phac-1(lf)* (Figure 6c). The effects of *snn-1(S9A)* on the thermotaxis learning phenotype appeared very similar: both *phac-1(gf)* and *snn-1(S9A)* mutants moved toward 22°C, but *snn-1(S9A)* had no effects on the cryophilic phenotype of *phac-1(lf)* raised at 24°C. Therefore, although the genetic interactions suggest that PHAC-1 and SNN-1 activities converge on synaptic functions, we cannot conclude that regulation of SNN-1 phosphorylation at Ser9 accounts for the effects of PP1-PHAC-1 signalling at synapses (Figure 6d).

**Figure 6.**
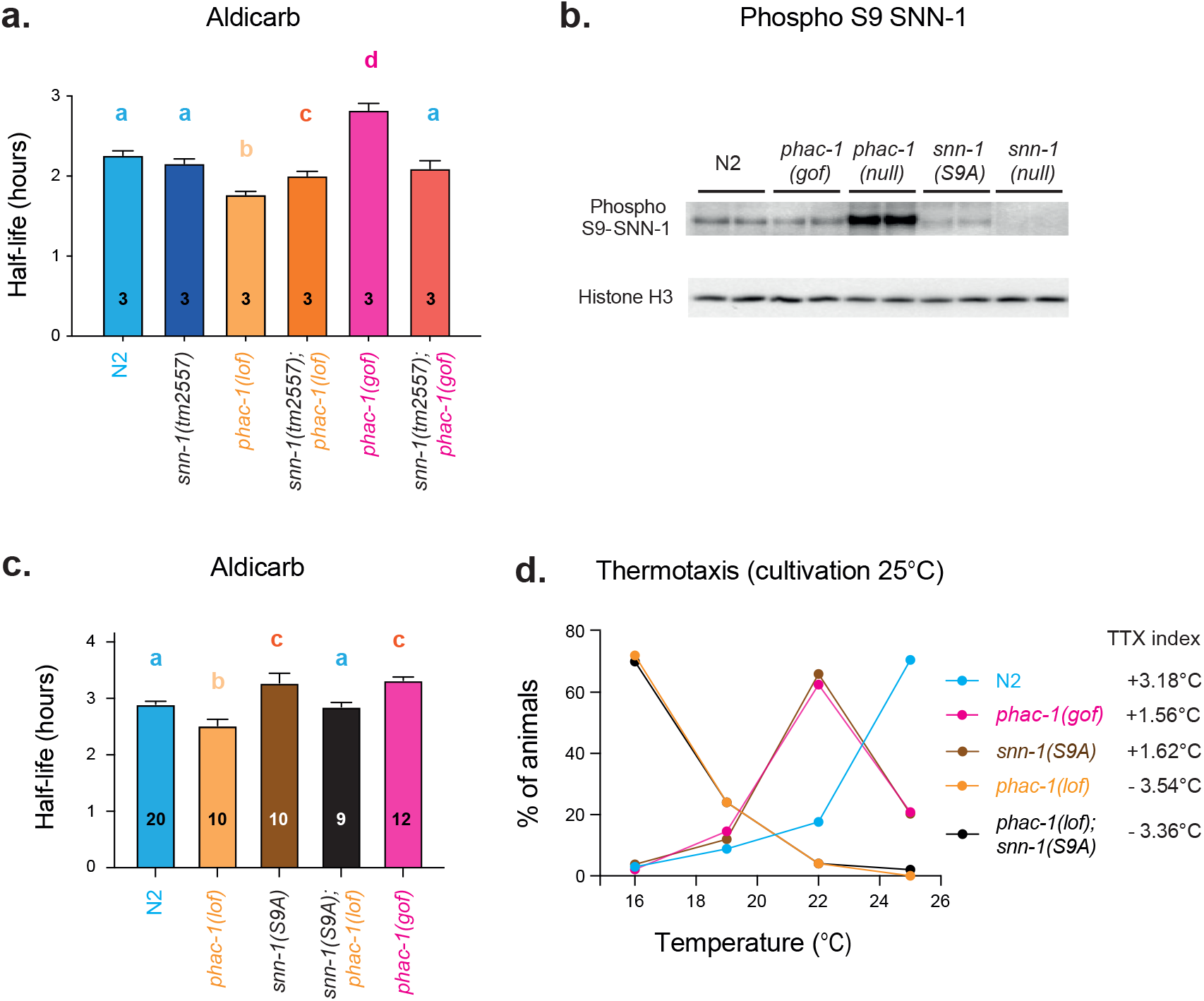
*phac-1* and *snn-1* act together to regulate the NMJ. **a)** The Aldicarb assay is used for epistatic analysis of the interactions of *phac-1(lf) and (gf)* alleles with *snn-1(tm2557).* The intermediate Aldicarb phenotypes of the double mutants *phac-1(lf);snn-1(tm2557) or phac-1(gf);snn-1(tm2557)* double mutants suggest but do not prove that *snn-1* mediate the Aldicarb phenotype of *phac-1* mutants. **b)** Western blots were performed using an antibody against the phosphorylated Ser9 of SNN-1. Histone H3 provides a loading control. The antibody specificity was demonstrated in *snn-1(tm2557)* null mutants and in the phosphodeficient *snn-1(S9A)* animals. SNN-1 S9 phosphorylation is increased in *phac-1(lf)* and weakly reduced in *phac-1(gf)* animals (quantification in Supplementary Figure 4f). **c,d)** Epistasis analysis of *phac-1(lf) and (gf)* alleles with the synapsin phosphodeficient allele *snn-1(S9A)* analysed using the Aldicarb **(c)** and thermotaxis assays **(d).** Although *snn-1(S9A)* phenocopies *phac-1(gf),* the Aldicarb phenotype of the *phac-1(lf);snn-1(S9A)* double mutants suggests that *snn-1* is not the only effector of *phac-1*. **d)** The thermotaxis assay used animals habituated to 25°C on food for 24 hrs prior to the assay. *phac-1(gf)* and *snn-1(S9A)* shows similar weakly cryophilic behaviour. However, the cryophilic behaviour of *phac-1(lf)* animals is not modified in *phac-1(lf);snn-1(S9A)* double mutants. Thermotaxis indexes are indicated. Data are shown as plotted as the mean ± SEM. For 6a and 6c, significant differences in pairwise comparisons are indicated by compact letters display. In brief, genotypes that share the same letter are statistically indistinguishable to each other, genotypes that have a different letter statistically differ (P<0.05) using One-way ANOVA and *Tukey’s* multiple comparison test.

## Discussion

Modulation of locomotor circuitry by neuropeptides was previously demonstrated to regulate E/I imbalance in the *acr-2(gf)* model [13]. We have used forward genetics to identify genes involved in NMJ neuromodulation by overexpressing the neuropeptide FLP-18, which perturbs E/I balance and confers striking behavioural phenotypes. We find that disrupting the sole *C. elegans* orthologs of PHACTRs and Synapsins, respectively called *phac-1* and *snn-1*, suppressed these defects. Variants of PHACTR1 and Synapsin I can cause human epilepsy, underscoring presynaptic mechanisms in controlling E/I imbalance [29, 36, 68]. While *prprp-1(lf)* prevented E/I imbalance and convulsions in the *acr-2(gf)* model, *phac-1(gf)* caused E/I imbalance and convulsions in the presence of the pro-convulsive drug PTZ.

We provide epistatic evidence that *phac-1* regulates signalling at the adult cholinergic NMJ in parallel and/or upstream to *snn-1.* Synapsin is one of the most abundant phosphoproteins associated with SVs and plays a critical role in SV dynamics [69]. Interestingly, Ser9 SNN-1 phosphorylation is regulated by PHAC-1 signalling. We investigated the possibility that direct dephosphorylation of the conserved Serine 9 of SNN-1 by the PP1-PHAC-1 holophosphatase might mediate some of the presynaptic effects of PHAC-1 signalling. Although the phosphodeficient allele *snn-1(S9A)* reduces cholinergic signalling at the NMJ it does not supress the effects of *phac-1(lf),* suggesting a more complex picture than this linear mechanism of action. This result either reflects multiple substrates of PHAC-1 at these synapses, and/or suggests SNN-1 is indirectly dephosphorylated by PP1-PHAC-1 signalling. Importantly, the consensus motif previously established for PHACTR1 substrates does not match the sequence surrounding the conserved Ser9 residue in SNN-1 and Synapsin I [34].

Multiple *de novo* mutations have been identified that cause autosomal dominant Developmental Encephalopathy and Epilepsy (referred to as DEEs) in infants. These include variants of genes involved in presynaptic function such as STXBP1/UNC-18 [70, 71]. All variants associated with the DEE70 subgroup of conditions replace highly conserved amino acids of PHACTR1 within motifs involved in G-actin binding [36]. A previous publication combining knock-down of PHACTR1 with the expression of rescue constructs in the embryonic mouse brain suggested that 4 DEE70 variants had dominant negative effects: N479I, L500P, I518N, R521C [36]. Invertebrates in which the neurotransmitter release machinery is highly conserved provide accessible model organisms to study the mechanism of action of variants causing synaptopathies [72, 73]. Here, we characterized *C. elegans* strains lacking *phac-1* or mimicking the DEE70 variants in the endogenous *phac-1* locus. We show that *L500P*, *L519R*, and *R521C* mutations in PHAC-1 associated with DEE70, cause a gain-of-function phenotype in *C. elegans* resulting in lethargic locomotion and impaired Aldicarb-sensitivity at the NMJ, while loss of *phac-1* induces the opposite phenotype. PHACTRs are though to cycle between an active holophosphatase state when bound to PP1, and an inactive, G-actin bound state [48–50]. Reducing G-actin binding to PHAC-1 phenocopied the Aldicarb-resistance of DEE70 variants, while reducing PP1 binding to PHAC-1 phenocopied the Aldicarb-sensitivity of *phac-1(lf),* suggesting DEE70 variants reduce G-actin binding to PHAC-1 and favour PP1 binding. Consistent with this, fusing PP1 to PHAC-1 phenocopied the Aldicarb-resistance of DEE70 variants. We therefore conclude that DEE70 variants promote the active state of the holophosphatase. These results contrast with the dominant negative effects observed in [36], but confirm the effect predicted by [35, 48–50]. In *C. elegans*, the hyperactive PP1-PHAC-1 holophosphatase acts post-developmentally to regulate neuromuscular function. As we obtain a partial rescue of *phac-1(lf)* by neuronal expression of human PHACTR1, it is likely that some of the substrates and functions of the PP1-PHACTR1 holophosphatase are conserved in neurons. Our results confirm the pathogenicity of DEE70 missense variants in *C. elegans,* provide evidence of its dominant positive effects, and determine one potential mode of action for DEE70 in neurons: PHAC-1 signalling controls the presynaptic SVs and DCVs pools, suggesting a presynaptic mode of action.

Several lines of evidence suggest that *phac-1(gf)* mutants have reduced or delayed SV recycling: 1) The total number of synaptic SVs assessed by quantitative imaging and by EM is reduced in DA presynaptic terminals; 2) SVs have increased diameter, supporting a defect in SV recycling; 3) SNB-1-pHluorin fluorescence is increased, indicating SV release, retrieval and/or reacidification are impacted; 4) reduced synaptic content of two endocytic markers, ITSN-1-GFP and UNC-57-GFP, are consistent with endocytic defects. Several observations suggest opposite alterations of NMJ signalling in *phac-1(lf)* s compared to *phac-1(gf)* mutants. These include the reduced locomotory behavior and Aldicarb-hypersensitivity of *phac-1(lf)* animals. Although Aldicarb-hypersensitivity suggests increased cholinergic signalling at the NMJ, we did not observe evidence for differences in cholinergic excitability or SV exocytosis properties in *phac-1(lf)* and *phac-1(gf).* Instead, both *phac-1(lf)* and *(gf)* mutants present evidence of SV recycling defects, such as larger SV diameters, increased fluorescence of the SNB-1-pHluorin or the expanded presynaptic terminal of hypersecretory *phac-1(lf)*. It is therefore unlikely that SV recycling defects alone would explain the opposite aldicarb-sensitivity phenotypes.

Our genetic screen suggests FLP-18 neuropeptide signalling is reduced in *phac-1(lf)* mutants. Although reduced neuropeptide secretion could not be demonstrated in *phac-1(lf)*, we did observe a significant accumulation of DCVs distal to dense projections in EM analysis, and ectopic DCVs in quantitative imaging of NLP-21-GFP, suggesting reduced *phac-1* signalling affects DCV biology. Supporting a role for PHAC-1 signalling in the control of neuropeptide secretion, we observed increased NLP-21-GFP secretion from DA cholinergic motorneurons of *phac-1(gf)* animals. Multiple feedback mechanisms employ neuropeptides, bioamines and extra-synaptic signalling to fine-tune the *C. elegans* locomotory circuit, and to adapt locomotion patterns [13, 74, 75]. Notably, neuropeptides processed by the peptide convertase EGL-3 compensate for the E/I imbalance caused by *acr-2(gf)* by increasing GABAergic transmission [13]. Although our results suggest *phac-1* primarily controls cholinergic motorneuron signalling, the hypersensitivity *phac-1(gf)* animals to PTZ suggests GABAergic signalling is also modulated by *phac-1*. We suggest that the PP1-PHAC-1 complex controls peptidergic release from cholinergic motorneurons and thereby modulates GABAergic NMJs to balance E/I.

PHAC-1 signalling varies according to actin polymerisation and local G-actin depletion [48–50]. Stable and dynamic actin pools are observed at presynaptic terminals [76], and actin polymerisation controls SV endocytosis, recycling and traffic [77–80]. PHAC-1 might act as a sensor of presynaptic depletion of G-actin. While increased PHAC-1 signalling disrupts E/I balance in the presence of PTZ, E/I imbalance is improved by reducing PHAC-1 signalling in *acr-2(gf)* mutants. Varying PP1-PHAC-1 signalling may survey presynaptic function and, by influencing peptidergic release and SV recycling, promote the E/I balance. Similarly, PHACTR1 signalling may be involved in tuning the excitatory/inhibitory (E/I) balance of mammalian cortical circuits.

## Methods

### Strains

Strain maintenance and genetic manipulation were performed as previously described [9]. Animals were cultivated at room temperature on nematode growth medium (NGM) agar plates seeded with OP50 bacteria. On the day before experiments, L4 larval stage animals were transferred to fresh plates seeded with OP50 bacteria. The strains generated or obtained, as well as the primers used for genotyping, are provided in Supp List 1

### Constructs, transgenes and germline transformation

*C. elegans* N2 genomic DNA or cDNA was used as template for cloning PCRs. Cloning PCRs were performed using Phusion High Fidelity DNA polymerase (M0530L, New England BioLabs) and then validated by Sanger sequencing. Details on primer used are provided in Supp List 1. Expression plasmids used to generate transgenic animals were generated by means of Multisite Three-Fragment Gateway Cloning (Invitrogen, Thermo Fisher Scientific, MA). To generate the PHAC-1(2R/A) plasmid, site-specific mutagenesis was performed by standard PCR using a plasmid containing wild-type *phac-1* cDNA sequence as template. Primers used are provided in Supp List 1. Details on Expression plasmids are provided in Supp List 1. All transgenic worms were generated by microinjection using standard techniques [81]. For most injected constructs, injection mixes were composed of 30 ng/μl targeting constructs, 30 ng/μl of co-injection markers, and 40 ng/μl of 1 kb Plus DNA mass ladder (Invitrogen, Thermo Fisher Scientific) as carrier DNA, for a final injection mix DNA concentration of 100 ng/μl.

### Gene editing

Gene editing by CRISPR Cas-9 was done by direct injection of Cas9 guide-RNA ribonucleoprotein (RNP) complexes into the syncytial gonad as previously described [82]. CRISPR-mediated homologous recombination with a DNA repair template is increased by mixing, melting, and reannealing template dsDNA. An injection mix was prepared with 30 pmol of Cas9, 90 pmol of tracrRNA, 95 pmol crRNA, 500 ng melted dsDNA, 800 ng myo-2p::RFP plasmid. For *phac-1* gene editing, ∼20 injected P0 animals were cultured individually. We selected 3 P0 plates with ∼40% of the progeny carrying the myo-2p::RFP transgene to pick F1. We singled 24 F1 for point mutants and 48 for wrmScarlet insertion. *phac-1* gene editing was screened using PCR-restriction digest. The gRNA and repair template used are provided in Supp List 1.

### RNAi knockdown of Ex[phac-1::SL2-tagRFP]

Worms were raised on NGM plates containing IPTG and seeded with the *E. coli* HT115(DE3) strain transformed with the plasmid L4440 containing RNAi against F26H9.2/*phac-1*, against *dpy-11* (as positive control) or the empty vector as negative control. The F26H9.2 RNAi strain carries a 1194 bp PCR fragment between primers ATCACGACGGGCTACTTTTG and CGGATCAGCGTAAATTTGAA. As IPTG is light-sensitive, these strains were grown in the dark under otherwise normal conditions. Imaging was performed as described below.

### Morphological and behavioural video recording

Video recordings were acquired at 15 frames per second (fps) with 3 array cameras connected to a computer equipped with Motif Software (Loopbio, Wien, Austria). Worms were segmented, tracked and skeletonized using Tierpsy Tracker [83], freely available on https://github.com/Tierpsy. After analysis, the features of interest were extracted. For locomotion assays, 30 worms were placed on low peptone NGM with OP50 seeded ∼16hrs before. Each genotype was recorded 3 times. Worms were allowed to adapt to 7% O_2_ for 5 minutes before videorecording started. Worms were recorded for 5 minutes until the gas mixture pumped into the chamber was changed from 7% to 21% O_2_. Midbody forward speed parameter was extracted from the videos and compared across the different genotypes.

### Isolation of *flp-18(XS)* suppressors and identification

*flp-18(XS)* suppressors were identified in a forward genetic screen. Animals from the strain OQ1 (Is[pflp-18::flp-18-SL2-GFP; ccGFP] V; Key Resources Table) carrying stably integrated *flp-18(XS)* were mutagenized with EMS using standard procedures, and 8–12 P0 animals were isolated on 8 P0 plates [9]. Subsequently, 4–6 F1 animals from each P0 plate were placed on 15 NGM growth plates, and their F2 progeny examined for locomotion. 48 F2 mutants were isolated in which locomotion appeared restored, and worms from their F3 plates were scored. We scored from −1 to 4 (0 being unchanged and 4 being like N2) the following parameters typical of the *flp-18(XS)* strain: slow locomotion on food, and off food, deep body curvature during forward locomotion, frequent reversals, abnormally long reversals when prodded, long reversals with extreme curvature when prodded. Genomic DNA was isolated from the 25 best suppressor strains identified in the forward genetic screen. Genomics core facility of ULB performed paired end 150 bp sequencing on Illumina MiSeq; 20X coverage. Sequences were aligned to the original OQ1 and N2 strains. All high score mutations within ORFs were highlighted, and candidate suppressors were ranked according to repeated observations within the 25 strains sequenced. We observed 3 mutations for *npr-5*, 3 mutations for *cmk-1*, 2 mutations for *crh-1,* 2 mutations for *ric-19.* Deletion appeared only once in *snn-1* and *phac-1*. The *ulb03 and ulb14* alleles were crossed to the polymorphic Hawaiian CB4856 strain. ∼60 F2 progeny of this cross all exhibiting GFP expression from the *flp-18(XS)* integrated construct but displaying improved locomotion (homozygous mutants for suppressor) were identified using a compound fluorescence microscope. Genomic DNA was isolated from their pooled progeny and subjected to sequencing. The density of unique and polymorphic single nucleotide polymorphisms was assessed using online tools (CloudMap) on the usegalaxy.org server as previously described [84]. The generated chromosomal level frequency plots for parental alleles showed a peak of unique variants on chromosome 1 for *ulb03* and 4 for *ulb14*. Within the middle of these peaks deletion mutations were identified for *phac-1* (*ulb03*), and *snn-1(ulb-14).* The alleles were outcrossed twice in N2 (CGC) before further analysis

### Aldicarb Paralysis Assay

Aliquots of stock Aldicarb (33386-SIGMA, 100mM in 70% (v/v) ethanol) were kept at 4°C until its use. Aldicarb plates (0.5mM or 1mM in low peptone NGM) were prepared at least three days before the experiments. Figure to figure variability in paralysis dynamics reflect variability in aldicarb concentration from stock and/or in batches of plates. As much as possible, all replicates of each figure were collected with a single batch of plates, for all genotypes. L4 worms were picked 20 hours before the assay. 1 hour before the assay, 10μl of OP50 were placed carefully on each Aldicarb plate to maintain the population of animals in the center of the plate. Animals were placed on the bacterial lawn and paralysis of each worm was tested every 30min. The criterion for paralysis was the absence of midbody movement after poking three times with our worm picker. Paralyzed worms were removed from the plates and recorded. For each strain, the experiments were performed 3 to 14 times with 20 to 30 animals per replicate. For each replicate, the half-life of these paralysis curves was calculated by a fitting curved generated by the Sigmoidal curve fitting equation build in GraphPad (8.4) Prism (San Diego, CA). The presented data are the mean paralysis half-life for 3 to 14 replicates ± SEM. As controls we used *egl-21*(n476) and N2.

### Levamisole assays

Levamisole aliquots (L0380000-MERCK) were freshly prepared before the assay. The assays were performed essentially as described [85]. Briefly, low peptone NGM agar plates supplemented with levamisole at concentrations ranging from 0.2 to 0.8mM were prepared. L4 larvae were isolated 20hrs before the assay. For each plate, worms were placed onto the levamisole plates and inspected for induced paralysis every 10 minutes. Each plate corresponds to 20-25 adult worms, we made 3 replicates for each genotype, and the experiment was performed 3 times. Worms were considered paralyzed when they did not respond to three consecutive proddings with the platinum wire pick.

### PTZ assays

NMG plates containing 10mg/ml final concentration of PTZ (P6500-MERCK) were prepared the day before the assay. 20-30 one day old adult worms of each genotype were placed on the PTZ plates. After 30 min, we counted the percentage of worms that were showing head convolution also called ‘head bobs’. For every group 3 replicates were performed, and the experiment was repeated 3 times.

### Generation of (phosphoS9)-SNN-1 antibody and Western blot

The antibody against phosphorylated SNN-1 was raised against the peptide sequence FKRKF-S(PO3H2)-FSEDEG with KLH carrier protein by Eurogentech. Western blotting was performed as described in [86]. Worms were washed thrice with M9 buffer and re-suspended in phosphate buffer saline (PBS). These worms were sonicated in PBS containing a protease inhibitor cocktail and a phosphatase inhibitor. Sonication was performed at 25 amplitudes for 3 min with a pulse time of 15 seconds. The sonicated lysate was centrifuged at 16,000X g for 30 min at 4°C to remove debris, and supernatant containing the total protein was kept. 30µg of total protein was loaded in 10 % sodium dodecyl sulfate polyacrylamide gel electrophoresis (SDS-PAGE). The primary antibodies used for Western blot were anti-phosphorylated SNN-1 (1:1000) and anti-histone H3 antibody (1:1000). Chemiluminescence was detected using LAS 3000 GE ImageQuent and densitometric analysis was performed by ImageJ software (Image J, National Institute of health, Bethesda, MD).

### Thermotaxis assay

We collected eggs in a defined time window so as to culture around 100 worms, and grew each group at 16°C or 25°C on NGM plates with OP50 food. Adult worms were collected and washed once in M9. We set up a temperature gradient on an NGM square plate without bacteria and placed around 100 worms in the middle of the plate, the excess buffer was removed. After one hour we counted the number of worms present at each section of the plate corresponding to the different temperatures.

### Fluorescence imaging

Worms were synchronized either by bleaching a population of gravid adults, or using an egg-laying window. Worms were reared at 20°C and imaged at day 1 adulthood unless otherwise stated. Worms were mounted on 2% agarose pads diluted in M9 solution, placing them in a droplet of M9 on the agar pad and then cover them with a cover slip. Animals were immobilized with 10mM Levamisole (L9756-Sigma) diluted in the agar pad and in the M9 droplets. Levamisole agar pads and M9 dilutions were freshly made for each strain. Levamisole is a nicotinic acetylcholine muscle receptor agonist that induces paralysis after ∼10 minutes.

Images were acquired at the Light Microscopy Facility LiMiF (http://limif.ulb.ac.be) on a LSM780NLO confocal system fitted on an Observer Z1 inverted microscope (Carl Zeiss, Oberkochen, Germany). Images were acquired using a LD C Apochromat 40×/1.1 W Korr M27 objective or an alpha Plan Apochromat 63x/1.46 Oil Korr M27 objective. The settings were as follows: frame size was set at 1024 × 1024 pixels with a pixel size of 0.13 μm × 0.13 μm, pinhole size was set to 1 Airy Unit, Z-step optical sections were acquired at 0.4 μm steps. For individual channel acquisitions, the Main Beam Splitter matched the excitation wavelength of each used fluorophore. The following fluorophores excitation (Ex) and detection wavelengths (DW) were used: mEGFP (Ex: 488 nm – DW: 493–569 nm) and wrmScarlet/mKate/TagRFP (Ex: 543 nm – DW: 570–695 nm). Fluorophore excitation and detection wavelength ranges were set according to the information available at the FPbase database for each fluorophore [87]. Laser power and detector gain settings were adjusted to maximize signal-to-noise ratio and minimize saturation. Images were saved in .czi Zeiss file format. For examination of different genetic backgrounds, each strain was imaged on at least 2 independent days (always including control N2 and tested mutants), and the data were pooled together. Acquisition settings were the same across session and across genotypes per presynaptic marker for quantitative analysis.

### Image processing

Confocal images were processed using FIJI [88] and Zen 2.6 Pro (Blue edition) software (Carl Zeiss). Z-stack acquisitions were converted into a 2D image using maximum intensity projections to obtain a flattened image representative of the 3D volume. For fluorescence along the DA axon: we imaged a 100μm x 3μm area along the dorsal cord right after the posterior pharynx bulb and corresponding mostly to the DA1 axon. A Z-stack was obtained for each worm at 0.4 μm steps. A line was traced along the axon signal to collect fluorescence intensity along the axon. The same line was slightly shifted away from the axon to collect background fluorescence. Fluorescence intensity along the axon minus background fluorescence was analyzed using custom software written in Igor Pro (Wavemetrics), as described previously [89]. Peak fluorescence was detected if 2x brighter than the median plus 20% standard deviation, and width was >0.3, <10 μm at half-maximum. These bright puncta corresponded to local accumulation of the marker and were further analysed to extract their fluorescence intensity, density per μm.

For AIY presynapse and for coelomocyte fluorescence quantification: ROIs were drawn and used to quantify the fluorescence of the synaptic area or the coelomocyte as well as the background fluorescence. Area fluorescence (CTFC) was extracted using the following formula: CTFC = Integrated Density – (Selected Area × Mean fluorescence of background readings).

### Electrophysiology

Worms were immobilized on a sylgard-coated coverslip using Histoacryl Blue glue, and a lateral cuticle incision was made with a glass needle, exposing the ventral medial body wall muscles. Nematodes were viewed during recordings using a 40X water immersion objective on a Zeiss Axioskop. Body wall muscle recordings were made in the whole-cell voltage-clamp configuration (holding potential, −60mV) using an EPC-10 patch-clamp amplifier (HEKA, Lambrecht, Germany) and digitized at 1kHz. Voltage-clamp recordings were typically maintained for 2 - 5 minutes. Voltage clamping and data acquisition were controlled by Pulse software (HEKA) run on a Dell computer. Modified *Ascaris* Ringer contained (150mM NaCl, 5mM KCl, 5mM CaCl_2_, 4mM MgCl_2_, 10mM glucose, 5mM sucrose, and 15mM HEPES (pH 7.3, ∼340mOs). Fire-polished recording pipets with resistances ranging from 5-9 MΩ were pulled from borosilicate glass (World Precision Instruments). The patch pipette was filled with 120mM KCl, 20mM KOH, 4mM MgCl_2_, 5mM(N-tris[Hydroxymethyl] methyl-2-aminoethane-sulfonic acid), 0.25mM CaCl_2_, 4mM Na_2_ATP, 36mM sucrose, and 5mM EGTA(pH 7.2, ∼315mOsm). Data were acquired using Pulse and Patchmaster software (HEKA, Southboro, Massachusetts, United States). Subsequent analysis and graphing was performed using Pulsefit (HEKA), Mini analysis (Synaptodoft Inc, Decatur, Georgia, United States) and Igor Pro (Wavemetrics, Lake Oswego, Oregon, United States). Stimulus-evoked EPSCs were generated by placing a borosilicate pipette on the ventral nerve cord (one muscle distance from the recording pipette) and applying a 0.2 ms, 20V square pulse using a stimulus current generator (WPI). Evoked train data were acquired at a frequency of 20Hz (50 ms interpulse interval) for 10 pulses.

### Electron Microscopy

The worm strains were prepared using high pressure freeze fixation and freeze substitution (HPF/FS) as previously described [90, 91]. Briefly, twenty to thirty young adult worms were placed in specimen chambers filled with *E. coli* and frozen at −180°C, using liquid nitrogen under high pressure (Leica HPM 100). After freezing, samples underwent freeze substitution (Reichert AFS, Leica, Oberkochen, Germany) using the following program: −90°C for 107 hours with 0.1% tannic acid followed by 2% OsO_4_ in anhydrous acetone, incrementally warmed at a rate of 5°C/hour to −20°C, and kept at −20°C for 14 hours before increasing temperature by 10°C/hour to 20°C; samples were then infiltrated with 50% Epon/acetone for 4 hours, 90% Epon/acetone for 18 hours, and 100% Epon for 5 hours; finally, samples were embedded in Epon and incubated for 48 hours at 60°C [91]. Ultra-thin (40 nm) serial sections were acquired using an Ultracut 6 (Leica) and collected on formvar-covered, carbon-coated copper grids (EMS, FCF2010-Cu). Sections were post-stained with 2.5% aqueous uranyl acetate for 4 minutes, followed by Reynolds lead citrate for 2 minutes [91]. Images were obtained using a JEOL JEM-1400F transmission electron microscope, operating at 80 kV. Micrographs were acquired using a BioSprint 12M-B CCD Camera with AMT software (Version 7.01). Cholinergic synapses at the NMJ of the ventral nerve cord were identified based on established synaptic morphology [7]. Sections containing a DP, as well as two flanking sections on either side of the DP, were aligned and analysed blinded to genotype using TrakEM2 and ROI features of NIH FIJI/ImageJ software respectively [88, 92]. Synaptic vesicles were counted as docked when the vesicle membrane was fully contacting the plasma membrane of the neuron terminal (distance = 0 nm), vesicles that were within 1–5 nm of the plasma membrane that exhibited small tethers were not scored as docked. The distribution of docked vesicles from the DP was calculated for each section containing a DP, as well as one section on either side, using the ROI data from FIJI with Matlab scripts written by the Jorgensen Lab [93]. Values were imported to Prism (GraphPad) for graphing and statistical analysis using One-way ANOVA with Tukey post hoc analysis, or Kruskal-Wallis with Dunn’s test, for multiple comparisons.

### Data acquisition and statistical analysis

Each set of data represents the mean ± SEM of an indicated number (N) of replicates (for drug based paralysis assay) or animals (for all other experiments). Statistical significance was determined using Student’s-t test or one-way ANOVA followed by the appropriate test to control for multiple comparisons indicated in legends. Statistically, values were: >0.05 not significant (NS), *p≤0.05, **p≤0.01, ***p≤0.001. GraphPad Prism 8.4(San Diego, CA) was used to perform statistical tests and calculate P-values. In figures where several genotypes are compared together, we present the statistical differences between strains using the compact letters display. Shortly, genotypes that share the same letter are statistically indistinguishable to each other, while genotypes that have a different letter statistically differ (p<0.05).

## Acknowledgments

P.L. is a research associate of the Belgian National Fund for Scientific Research (FRS-FNRS). K.S., M.S-G., S.S. and P.L. are supported by grants from the FRS-FNRS. This work was supported by an Advanced ERC grant (269058 ACMO) to M.d.B. We thank the team of Alexander Gottschalk for the *snn-1(S9A)* strain. We thank the Imaging Facility of the Faculty of Medicine (LiMiF) of the ULB, supported by FRS-FNRS. This work made use of instruments in the Electron Microscopy Core of UIC’s Research Resources Center; and of the BioCryo facility of Northwestern University’s NUANCE Center, which has received support from the SHyNE Resource (NSF ECCS-2025633), the IIN, and Northwestern’s MRSEC program (NSF DMR-1720139). Some strains were provided by the CGC, which is funded by NIH Office of Research Infrastructure Programs (P40 OD010440).

## Author contributions

Conceptualization: P.L., M.D.B.; Formal analysis: K.S., M.S.-G., S.S., M.K., J.E.R. and P.L.; Funding acquisition: K.S., M.S.-G., M.D.B. and P.L.; Investigation: K.S., M.S.-G, S.S., M.K., J.E.R and P.L.; Resources: J.E.R., M.D.B. and P.L.; Supervision: J.E.R and P.L.; Visualization: K.S., M.S.-G., S.S, M.K., J.E.R and P.L.; Writing – original draft: P.L.; Writing – review and editing: K.S., M.S.-G., S.S., M.K., M.D.B, J.E.R and P.L.

## Inclusion and diversity

We support inclusive, diverse, and equitable conduct of research.

---

**Supplementary Figure 1 a)** Distribution of velocities (um/sec) for a population of ∼60 N2 and *flp-18(XS)* worms placed at 1% O2. N2 animals spend most of their time in forward locomotion at speed > 100um/sec and up to >200 um/sec. In comparison *flp-18(XS)* animals rarely speed faster than 200um/sec and frequently reverse (negative velocities). **b)** The average midbody curvature is increased during forward locomotion of *flp-18(XS)* compared to N2, the average speed of forward locomotion is reduced in *flp-18(XS)* compared to N2. **c)** The average midbody curvature is increased during backward locomotion of *flp-18(XS)* compared to N2, the average velocity during backward locomotion is similar in *flp-18(XS)* and N2.

**Supplementary Figure 2 a)** Alignment of the PHAC-1 C-terminal sequence with the *Drosophila melanogaster* ortholog CG32264 and the human orthologs PHACTR1 and PHACTR3. The 3 RPEL motifs are underscored in green, the PP1-binding domain (PP1BD) is underscored in orange. The 2R/A, R536P and DEE70-associated mutations are indicated. The alignment shows a high degree of conservation in the last 2 RPEL motifs and the PP1BD, including the amino acids associated with human DEE70. **b)** Forward locomotion speed of *phac-1* mutants compared to N2 controls: *phac-1(lf)* animals display reduced speed while *phac-1(gf)* animals display increased speed compared to N2. Data are plotted as the mean ± SEM; one-way ANOVA. **c)** Expression in D1 adults of wrmScarlet in the CRISPR knock-in reporter strain carrying *wrmScarlet* c-terminally fused to *phac-1 ORF*. Expression is observed in the pharyngeal muscles, head muscles, excretory cells and the nerve ring**. d)** A second reporter strain was designed. It carry an extra-chromosomal array where SL2-TagRFP forming an operon regulated by a 6kb fragment of the *phac-1* locus. Below are shown DIC (left) and fluorescence (right) images of embryonic (top) and L1 (bottom) stages. Expression of TagRFP (red and grayscale on the left) is widespread, including expression in neurons at all developmental stages. The coelomocytes co-injection marker is observed in green. **e)** Panneuronal expression of N- or C-terminal GFP fused to *phac-1* (top) is observed in the cytoplasm and neurites of neurons but not in the nucleus. The N- or C-terminal placement of GFP does not alter the subcellular location of the protein. Panneuronal expression of a *gsp-1::phac-1::GFP* fusion construct (bottom) show similar distribution of the constitutive PP1-PHAC-1 holoenzyme and PHAC-1.

**Supplementary Figure 3 a)** Representative graph showing fluorescence of the active zone marker, UNC-10 along the dorsal nerve cord. Fluorescence peaks correspond to the enrichment of UNC-10 at presynaptic puncta. **b, c)** Fluorescence intensity **(b)** and peak density **(c)** of the active zone marker UNC-10-GFP in *phac-1* mutants compared to N2. We do not observe any significant differences in fluorescence intensity or peak density, indicating that the organisation and density of the active zones are not modified in *phac-1* mutants. **d)** Fluorescence intensity of the Dense Core Vesicle (DCV) marker NLP-21-GFP in the somas of DA neurons in *phac-1* mutants and N2 control. No significant changes are observed in the *phac-1* mutants compared to controls. **e)** Fluorescence intensity of the NLP-21-GFP peaks detected along the DA axon in *phac-1* mutants and N2 controls. Peak fluorescence intensity is decreased in *phac-1(gf)* compared to N2. **f)** A thermotaxis assay performed in N2, *ttx-3*, *phac-1(lf)*, *phac-1(gf)* and *snn-1(S9A)* mutants habituated to 16°C on food for 24 hrs prior to the assay. All strains display the expected cryophilic behaviour. **g**) The general morphology of the AIY neurite was preserved based on GFP-RAB-3 fluorescence.

**Supplementary Figure 4 a-c)** The ultrastructure of cholinergic motorneurons synapses was observed by electron-microscopy. **a)** The Dense Projection (DP) area is not altered in *phac-1* mutants compared to N2, but is significantly reduced in *pprp-1(gf)* compared to *pprp-1(lf).* **b)** The number of large vesicle is reduced in *pprp-1(lf)* compared to N2. **c)** the number of endosome is not modified in *pprp-1* mutants compared to N2 controls. **d, e)** Fluorescence of the SNB-1-GFP marker Fluorescence intensity of the axon **(d)** and peak density **(e)** are not modified in *phac-1* mutants compared to N2. f) Quantification of 2 independent Western blots performed using an antibody against the phosphorylated Ser9 of SNN-1. The values correspond to density of the phospho S9-SNN-1 bands, normalised to the density of H3 bands and normalised to N2 values. Phosphorylation is significantly increased in *phac-1(lf)* and decreased in *snn-1(null)* compared to N2. RM-one way ANOVA, N=2. *p≤0.05, **p≤0.01, ***p≤0.001. Data are plotted as the mean ± SEM.

