## Supplementary figures and images for "Neuroendocrine control of synaptic transmission by PHAC-1 in *C. elegans*"

### Supplementary Figures1-4

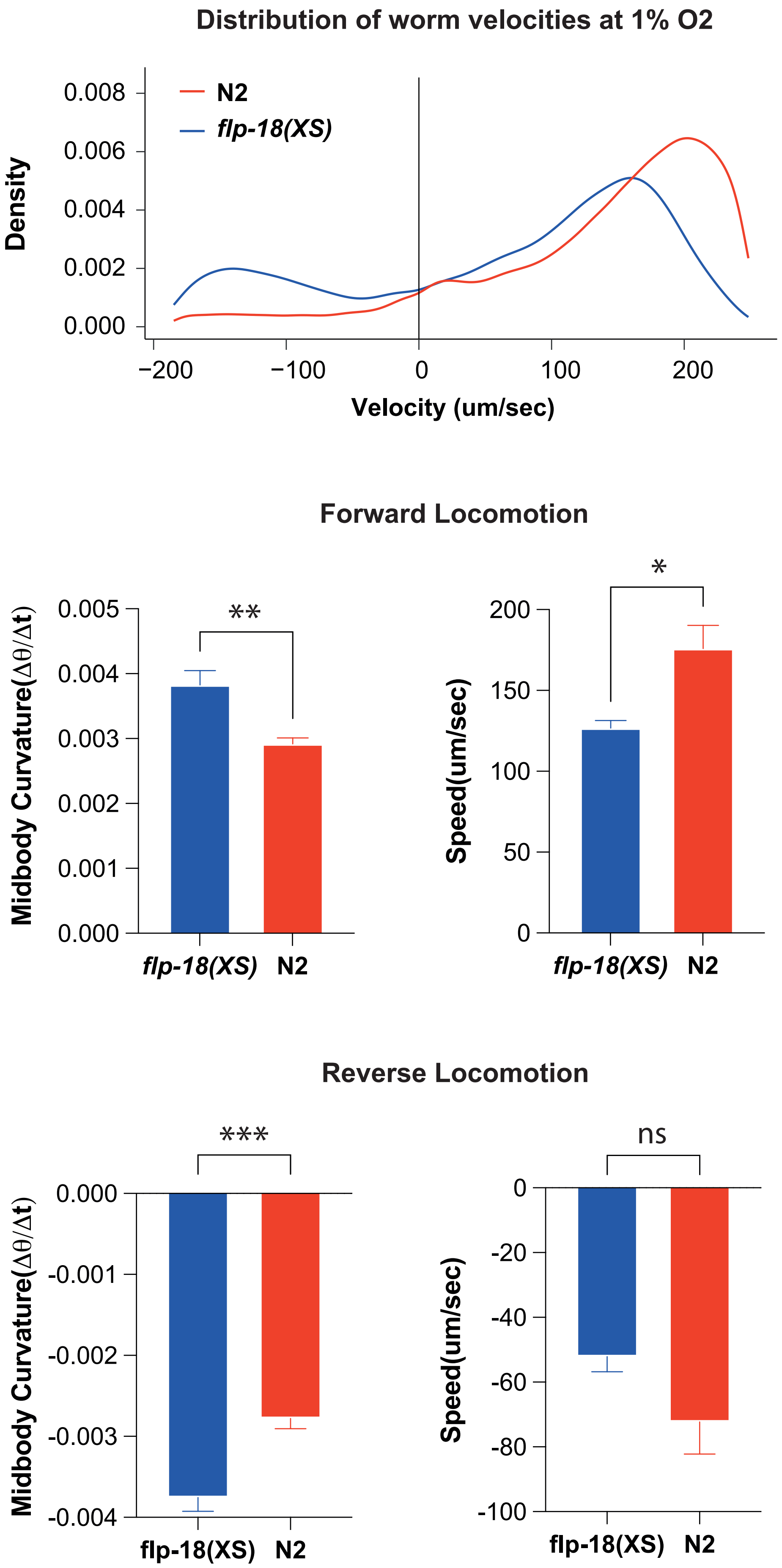

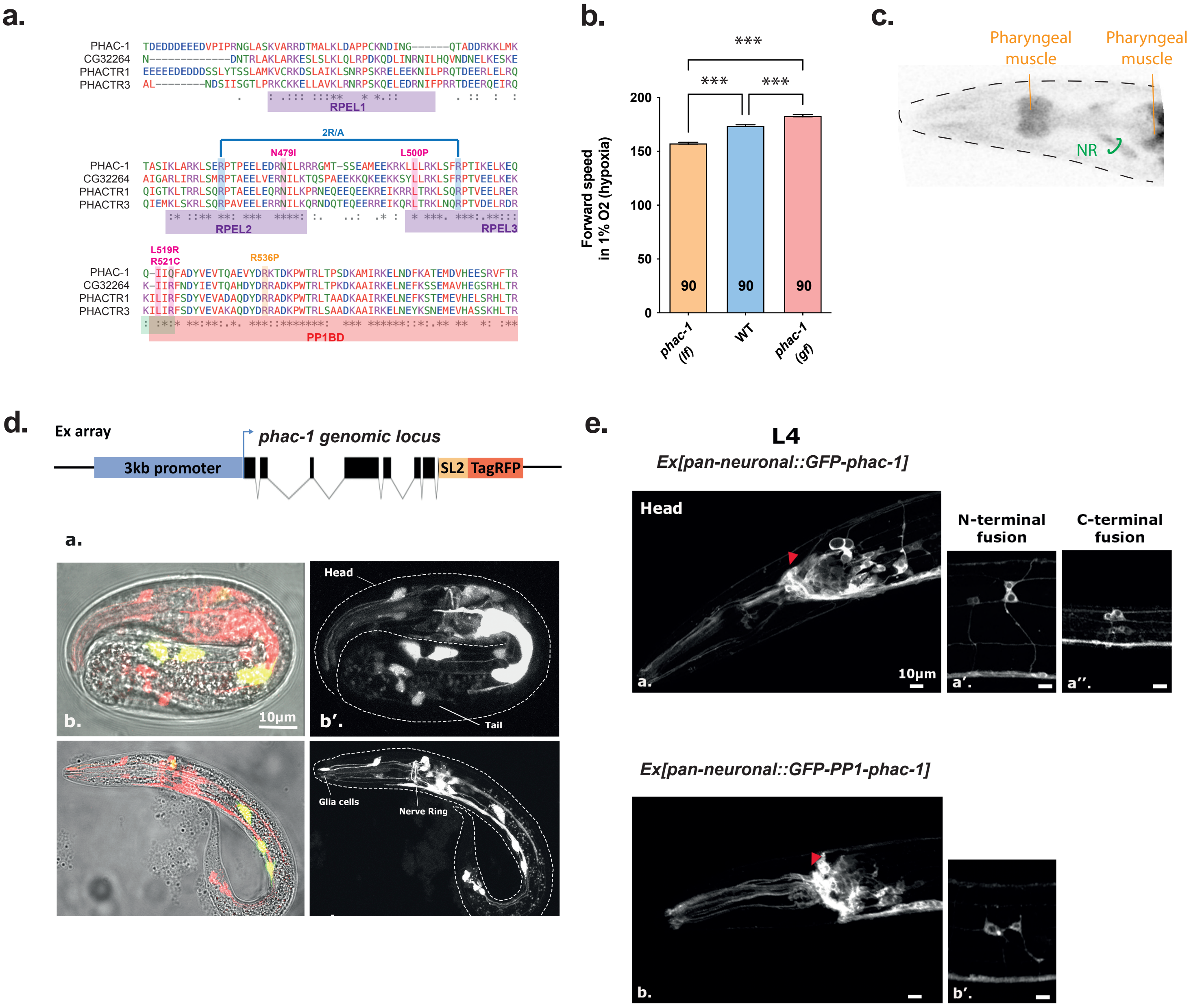

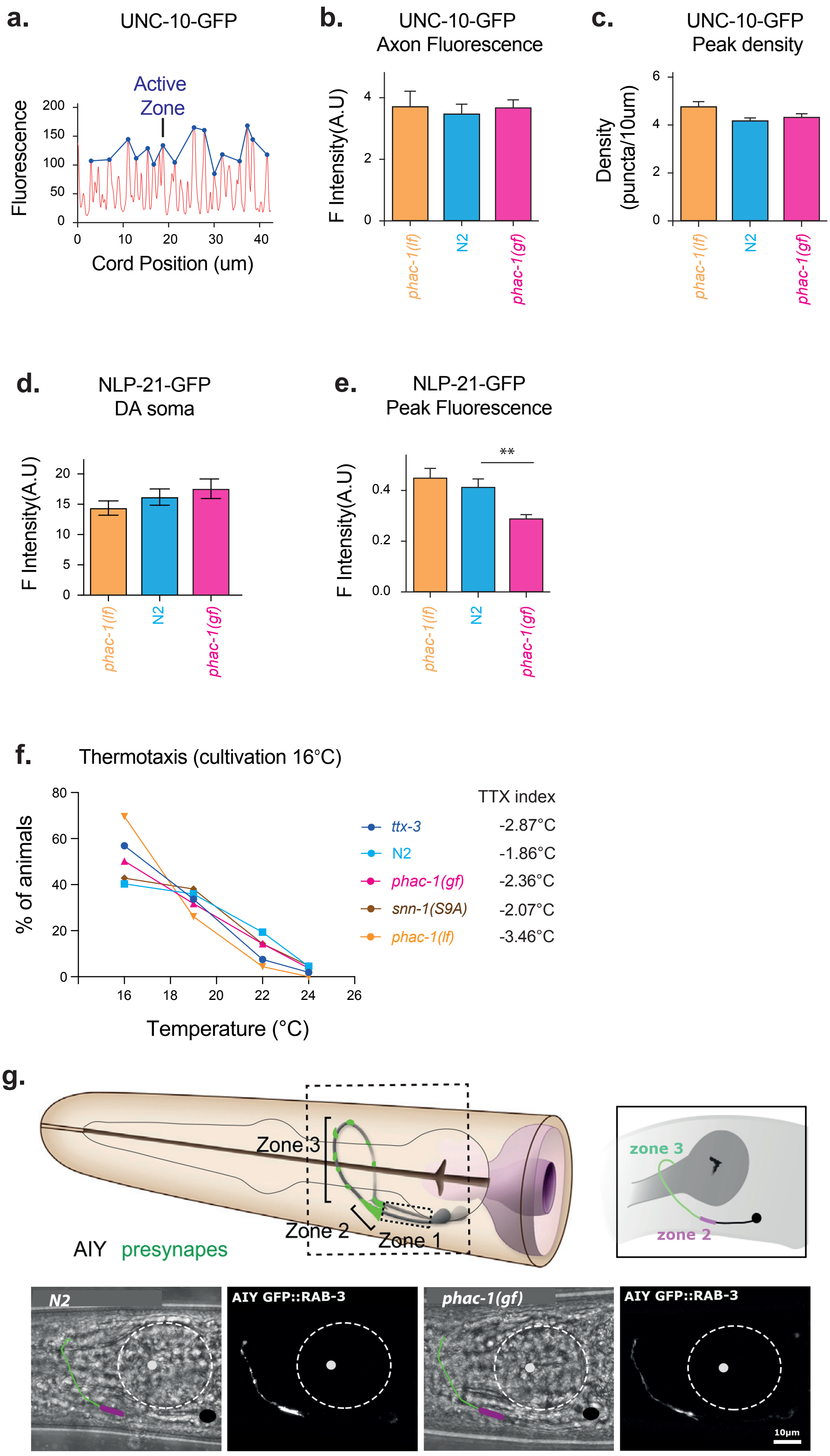

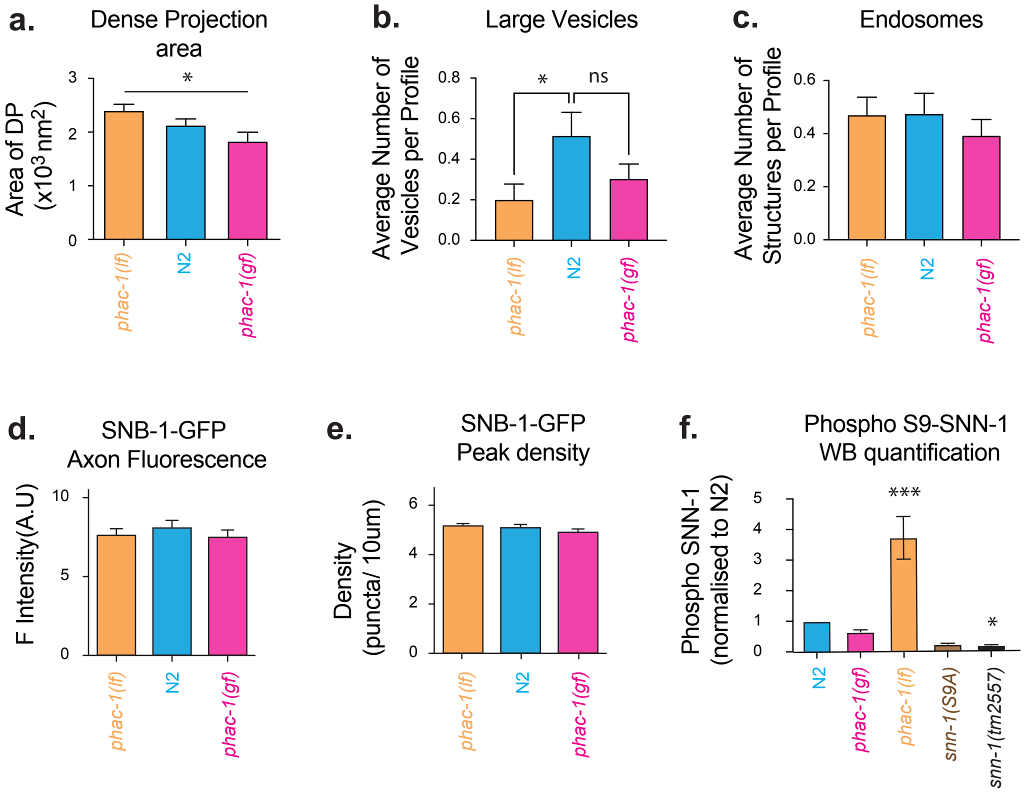
